# Molecular mechanisms of altered contraction with the β-myosin R403Q mutation in porcine ventricular muscle and a human stem cell-derived cardiomyocyte model

**DOI:** 10.1101/2025.08.27.672708

**Authors:** Sonette Steczina, Saffie Mohran, Matthew C. Childers, Timothy S. McMillen, Ateeqa Naim, Matvey Pilagov, Marica Dente, Kristina B. Kooiker, Christian Mandrycky, Khushi Tawde, Jennifer Hesson, Jing Zhao, Julie Mathieu, J. Manuel Pioner, Michael A. Geeves, Weikang Ma, Farid Moussavi-Harami, Neil M. Kad, Michael Regnier

## Abstract

The R403Q mutation in the sarcomere protein beta-myosin heavy chain (β-MHC) is a known genetic cause of hypertrophic cardiomyopathy (HCM), associated with ventricular hypercontractility, impaired relaxation, and cardiac arrhythmias. Despite extensive research, the mutations impact on myosin contractile properties remains unclear, likely due, at least in part, to discrepancies cross different model systems. In this study, we used a multidisciplinary approach to explore mutational effects using two distinct heterozygous R403Q systems: a Yucatan minipig model and human induced pluripotent stem cell-derived cardiomyocytes (hiPSC-CMs). X-ray diffraction of R403Q minipig ventricular muscle demonstrated reduced order of the thick filament, suggesting destabilization of the inhibited OFF (vs. ON) state of myosin in relaxed muscle, which correlated with elevated force at submaximal calcium. Super-resolution, single-molecule fluorescence microscopy indicated elevated ATPase activity in thick filament zones lacking cMyBP-C. Furthermore, R403Q myofibrils exhibited slower activation and relaxation kinetics, with reduced sensitivity to ADP. Molecular dynamics simulations suggested that altered interactions at the actomyosin interface contribute to these effects, rather than changes at the nucleotide binding pocket, typically associated with ADP release. Human engineered heterozygous R403Q hiPSC-CMs exhibited reduced maximal myofibril force, slowed contractile kinetics, and hypercontraction in engineered heart tissue constructs−consistent with HCM phenotypes observed in the heterozygous porcine model. Our results demonstrate that the R403Q mutation induces early and persistent contractile dysfunction, and that hypercontractility and slower contractile kinetics may result from a combination of an increased population of activated (ON) myosin heads and delayed detachment during cross-bridge cycling, respectively.

## II. INTRODUCTION

The first genetic mutation linked to hypertrophic cardiomyopathy (HCM) was the R403Q missense mutation in the sarcomeric protein beta-myosin heavy chain (β-MHC) (1). For patients with the R403Q mutation and many other HCM-causing mutations in sarcomere proteins, the phenotypic profile often includes ventricular hypercontractility, impaired relaxation, and a high incidence of cardiac arrhythmias (2). Due to the severity of the R403Q mutation, it has been a focus of several studies aimed at uncovering the pathological mechanisms that lead to disease progression.

There have been contradictory reports of the effects of the R403Q mutation on the kinetics of contraction and relaxation, likely due in part to the differences in model systems and myosin isoforms studied. Rodent hearts predominantly express the α-MHC isoform, and measures of myosin activity and product release rates associated with the cross-bridge cycle differ for α-MHC with the R403Q mutation when compared to transgenic mice expressing β-MHC with the R403Q mutation (3, 4). A more recent study using a heterozygous transgenic rabbit model, which expresses the β-MHC isoform, Lowey *et al*. (5) reported slower *in vitro* motility, unimpaired ATPase activity, and reduced force and velocities in isolated myofibrils with the heterozygous R403Q mutation. The contractile abnormality of isolated myofibrils with the R403Q mutation is inconsistent with results reported for myofibrils from human myectomy samples, which have consistently shown faster cross-bridge cycling kinetics for R403Q myofibrils compared to mutation negative control human myofibrils (6, 7). Furthermore, Witjas-Paalberends *et al*. proposed that the R403Q mutation significantly decreases myosin head efficiency, suggesting that myosin heads cycle faster and consume more ATP to generate a similar amount of force compared to controls (7). The faster myofibril cross-bridge cycling rates in human myectomy tissue is consistent with results from an earlier study on isolated β-MHC from human HCM R403Q patients, as determined by an *in vitro* motility assay (8). This study also reported a shorter power-stroke step duration, measured via single-molecule laser trap, which the authors postulated was due to altered ADP release or destabilization of strong actomyosin interactions. Other reports on ADP release have been contradictory and are also likely affected by the model system and myosin isoform used (4, 9), but destabilization of the actomyosin binding surface has been suggested from follow-up studies (9, 10).

The contradictory results among mouse models, transgenic rabbits, and human myectomy samples motivated the generation of a large animal model of the heterozygous R403Q mutation with a rapidly developing HCM phenotype of hypercontractility, hypertrophy, and an approximately equal ratio of wildtype and mutant protein (11–13). This model has been used as a tool to study the low ATPase, super-relaxed (SRX) state of β-MHC, the mode of action of the myosin targeted, small molecule inhibitor mavacamten (11), and for X-ray diffraction studies as a method to quantify cardiac thick filament structural disarray associated with HCM (14). However, the conflicting reports on contractile mechanics, kinetics, and energetics remain, and little is known about the altered myosin structure that underlies the contractile abnormalities. There is also a lack of information regarding when and how the disease process is initiated by the R403Q mutation.

To address these questions, we have employed a variety of biophysical, biochemical, and computational approaches in two model systems, the Yucatan minipig and human induced pluripotent stem cell-derived cardiomyocytes (hiPSC-CM) that contain the heterozygous R403Q mutation. Our results suggest that the R403Q mutation enhances myosin recruitment from the thick filament backbone, as seen by increased thick filament disorder in resting porcine tissue using X-ray diffraction. Single-molecule fluorescent ATP (Cy3-ATP) tracking using super-resolution microscopy revealed an increased population of high-ATPase-activity myosin heads relative to low-ATPase-activity myosin heads for R403Q myofibrils, specifically in the regions of the sarcomere where cMyBP-C is absent. This greater population of readily available myosin was correlated with enhanced force during sub-maximal calcium activation, consistent with calcium levels seen in cardiac function, manifesting as higher calcium sensitivity of myofibrils, cells, and tissue. Interestingly, maximal calcium activated force in demembranated ventricular tissue strips was slightly, but not significantly, reduced. Maximal specific force in isolated myofibrils was significantly reduced and the contraction and relaxation kinetics were slowed and less sensitive to elevated ADP. Molecular dynamics (MD) simulations of β-MHC bound to actin in the post-powerstroke, ADP-bound state suggest altered binding interactions of myosin at the actin interface, but no alteration in ADP nucleotide handling within the nucleotide binding pocket. In a human cell-based model of the R403Q mutation (heterozygous), myofibrils recapitulated results from adult minipig ventricle myofibrils, with reduced maximal force and slower relaxation kinetics. Twitch force measurements in engineered heart tissue (EHT) constructs demonstrated hypercontractility and delayed relaxation, consistent with HCM patient phenotypes. Together, our results provide increased insight into the multiple molecular mechanisms of hypercontractility and slowed relaxation resulting from the myosin R403Q mutation in cardiac muscle.

## III. METHODS AND MATERIALS

### Collection of porcine tissue

Isolated hearts from 1-month-old, sibling-matched WT (*N =* 3) and R403Q (*N =* 3) male Yucatan pigs were provided by Exemplar Genetics Inc. Humane euthanasia and tissue collection procedures were approved by the Institutional Animal Care and Use Committees at Exemplar Genetics. The procedures were conducted according to the principles in the “Guide for the Care and Use of Laboratory Animals”, Institute of Laboratory Animals Resources, Eighth Edition (15), the Animal Welfare Act as amended, and with accepted American Veterinary Medical Association (AVMA) guidelines at the time of the experiments (16).

### X-Ray diffraction

Left ventricular myocardium samples were prepared as described previously (17, 18). Briefly, samples of frozen left ventricular myocardium were permeabilized in relaxing solution (2.25 mM Na_2_ATP, 3.56 mM MgCl_2_, 7 mM EGTA, 15 mM sodium phosphocreatine, 91.2 mM potassium propionate, 20 mM imidazole, 0.165 mM CaCl_2_, 15 U/mL creatine phosphate kinase containing 15 mM 2,3-butanedione 2-monoxime (BDM), 1% Triton X-100, and 3% dextran at pH 7) at room temperature for 2 – 3 h. Muscles were then washed with fresh relaxing solution for ∼10 mins, 3 times. The tissue was dissected into ∼300 μm diameter fiber bundles and clipped with aluminum T-clips. The preparations were then stored in cold (4 °C) relaxing solution with 3% dextran for the day’s experiments. X-ray diffraction experiments were performed using the small-angle X-ray diffraction instrument on the BioCAT beamline 18ID at the Advanced Photon Source, Argonne National Laboratory (19). The X-ray beam energy was set to 12 keV (0.1033 nm wavelength) at an incident flux of ∼5 × 10^12^ photons per second in the full beam. The X-ray beam was focused to ∼250 × 250 μm at the sample position and to ∼150 × 30 µm at the detector. The specimen-to-detector distance was approximately 3.5 m. The muscles were incubated in a customized chamber, and experiments were performed at 28 – 30 °C. The muscles were stretched to a sarcomere length of 2.3 µm by monitoring the laser diffraction patterns from a helium-neon laser (633 nm). The X-ray patterns were collected in relaxing solutions with 100% ATP or 100% dATP on a MarCCD 165 detector (Rayonix Inc., Evanston, IL) with a 1 s exposure time.

To minimize radiation-induced muscle damage, the muscle samples were oscillated along their horizontal axes at a velocity of 1 – 2 mm/s. The irradiated areas were moved vertically after each exposure to avoid overlapping X-ray exposures. The data were analyzed using the MuscleX software package developed at BioCAT (20). The equatorial reflections were measured by the “Equator” routine in MuscleX to calculate the equatorial intensity ratio, I_1,1_/I_1,0_, and inter-filament lattice spacing, d_1,0_, as described previously (21). The intensities and spacings of meridional and layer line reflections were measured using the “Projection Traces” routine in MuscleX after the patterns were quadrant-folded and the diffuse scattering subtracted to improve the signal-to-noise ratio, as described previously (21, 22). Three to four patterns were collected under each condition, and spacings and intensities of X-ray reflections extracted from these patterns were averaged.

### Single-molecule ATP tracking

Porcine myofibril isolation, fluorescent staining, image acquisition, and data analysis for single-molecule ATP tracking were performed as recently published (23). Briefly, a ∼3 mm^2^ piece of –80 °C frozen porcine left ventricular tissue was cut from a larger segment of either WT or R403Q tissue and rapidly thawed in ice-cold Rigor Buffer (20 mM MOPS, 132 mM NaCl, 5 mM KCl, 4 mM MgCl_2_, 5 mM EGTA, 10 mM NaN_3_, 5 mM DTT, 20 mM BDM, protease inhibitor cocktail (A32965; Thermo Scientific), pH 7.1). While continuously submerged in Rigor Buffer, tissue was cut into 0.5 – 1 mm thick strips and pinned down at slack length with tungsten rods secured into a Sylgard™ polydimethylsiloxane (PDMS)-coated petri dish. Rigor Buffer was then substituted with a permeabilization solution (Rigor Buffer + 0.5% Triton X-100). Overnight permeabilization was performed with gentle agitation at 4 °C. The following day, permeabilization solution was replaced with fresh Rigor Buffer and washed three times. Tissue strips were then further cut down to 0.25 – 0.5 mm thick strips on ice submerged in Rigor Buffer and then transferred to a 2 mL micro-centrifuge tube with 500 μL of Rigor Buffer. To isolate single myofibril bundles, an ethanol-cleaned and water-washed Tissue Ruptor II was used at the lowest speed for 2 × 10 s with a 1 min rest on ice. The myofibril concentration for antibody staining was kept constant by diluting the isolated myofibril suspension to an optical density (OD) of ∼0.4 at 600 nm using a 7315 UV-VIS spectrophotometer (Jenway). To fluorescently label alpha-actinin containing Z-lines, myofibrils were simultaneously incubated with 12 – 18 nM anti-α-actinin mouse antibody (A7811, Sigma-Aldrich) and 6 nM Alexa-488 goat anti-mouse IgG (A11001, ThermoFisher Scientific) with 1 mg/mL bovine serum albumin (BSA, Sigma) at 4 °C for 1.5 h under constant rotation and protected from light.

During antibody incubation, myofibril imaging chambers were constructed as previously described (24). Briefly, ethanol cleaned, and plasma treated coverslips were coated with 5 μL of 15 μg/mL poly-L-Lysine (PLL, Sigma) and allowed to dry for 30 mins. Imaging chambers were subsequently assembled and 100 μL of Rigor buffer was flowed in prior to the entire myofibril-antibody incubation solution. This was incubated at 4 °C for 30 mins to allow for myofibrils to adhere to the coverslip surface and then washed twice with 100 μL of Wash Buffer (Rigor Buffer without BDM) with a 1 min pause at 4 °C in between washes. Prior to imaging, 100 μL of Imaging Buffer (Wash Buffer, 10 – 15 nM Cy3-ATP, 5 mM ATP, and 2 mM Trolox) was flowed into the imaging chamber. Cy3-ATP was produced and gifted by Dr. C.P. Toseland, University of Sheffield, Sheffield, UK (25). For dATP experiments, the 5 mM ATP in the above Imaging Buffer was replaced with 5 mM dATP (GenScript).

Microscope specifications and image acquisition scheme have been described in a recent publication (23). Briefly, an oblique angle fluorescence (OAF) microscope was used with a 561 nm laser (20 mW, OBIS LS laser, Coherent, USA) to excite Cy3-ATP and a 488 nm laser (20 mW, Oxxius) to excite the α-actinin labeled Z-lines and fluorescence images captured using an OrcaFlash 4.2 camera (Hamamatsu). Stroboscopic illumination was used with 5 sequential frames each consisting of 200 ms 561 nm laser exposure followed by 1.8 s with no illumination. The 6^th^ frame included an additional 200 ms frame exposure using the 488 nm laser during the 1.8 s dark period of that frame. This illumination pattern was repeated for 1000 frames and controlled by a custom-built Arduino system. One myofibril was imaged per chamber to minimize build-up of ADP.

As utilized by Pilagov *et al*., (23, 24) the sequence of 1000 frames per myofibril was split into two stacks, one containing all sequential Z-line frames and one containing all sequential Cy3-ATP frames. Any myofibril that exhibited drift in the x-or y-direction was corrected using the Drift Correct ImageJ plugin to ensure that Z-lines remained in the same location across the duration of the video. Then TrackMate, ImageJ plugin (26), was used to extract the spatiotemporal location of each Cy3-ATP molecule and Z-line within each myofibril. In tracking the binding lifetime of individual Cy3-ATP molecules, a linking and gap-closing distance of 100 nm per track was used as well as a 5-frame gap-closing max frame gap.

The spatial resolution of TrackMate’s results for each Cy3-ATP binding event within each myofibril were superimposed on the TrackMate determined spatial localization of the fluorescent Z-lines using Microsoft Excel. Each sarcomere was analyzed individually by first fitting two straight lines through two consecutive Z-lines, correcting any tilt in the Z-lines, and subsequently correcting the placement of the Cy3-ATP molecules with respect to the tilt corrected Z-line positions. The position of each Z-line was determined by binning the x and y coordinates of these corrected Cy3-ATP molecules and then fitting them to a Gaussian distribution (24). The spatial precision of Cy3-ATP localization was previously determined to be 29.2 nm in the x coordinate and 26.7 nm in the y coordinate for this system (24). Sarcomere length was determined from the distance between two consecutive Z-lines. Data was only collected from sarcomeres with a sarcomere length ≥1.7 μm. For sarcomeres that met this criterion, the M-line was assumed as the mid-point between the Z-lines, and the sub-sarcomere zones were calculated based on the M-line. For this study on the cardiac muscle sarcomere, the P-zone was designated as 0 – 159 nm from the M-line, the C-zone was 160 – 500 nm from the M-line, and the D-zone was 501 – 800 nm from the M-line (27, 28). All Cy3-ATP molecules localized within a sarcomere were then binned into these three zones based on their x and y position relative to the Z-lines, any outside this range were discarded.

The location of each Cy3-ATP binding event within a sarcomere was also accompanied by the respective duration of attachment extracted by TrackMate. Each frame interval was 2 s. All Cy3-ATP data per sarcomere was analyzed together and separately per sub-sarcomere zones. Data across the entire sarcomere was fit to the sum of three exponential fits using Excel’s Solver GRG nonlinear function as in Pilagov *et al.* (23, 24). Data for each zone of the three zones was then fit to the sum of three exponentials again using Excel’s Solver GRG nonlinear function. Both an amplitude and a rate constant, *k*, were calculated per exponential. Using the rate constant (*k*_DRX_) of the DRX population and the rate constant (*k*_fit_) and amplitude (AMP_fit_) fit to the SRX population, we calculated a corrected amplitude for the SRX population based on the following equation: AMP_CORR_ = AMP_fit_*(*k*_DRX_/*k*_fit_) (24).

### Demembranated tissue mechanics

Frozen porcine tissue sections were thawed and permeabilized overnight at 4 °C in 50:50 glycerol relaxing solution (100 mM KCl, 10 mM MOPS, 5 mM K_2_EGTA, 9 mM MgCl_2_ and 5 mM Na_2_ATP (pH 7.0 with KOH), 1% (v/v) Triton X-100, 1% protease inhibitor (Sigma P8340), and 50% (v/v) glycerol). The following day, permeabilized tissue was washed and stored in the same 50:50 glycerol relaxing solution, but without Triton X-100. Samples were stored at –20 °C for up to one week. Thin strips of permeabilized porcine tissue were dissected and mounted between a force transducer (Aurora Scientific, model 400A) and a motor (Aurora Scientific, model 315C) using aluminum T-clips (Aurora Scientific) (29). Sarcomere length (SL) was set to ∼ 2.3 μm. Experiments were conducted in physiological solution (pH 7.0) at 21 °C containing a range of pCa concentrations (−log_10_[Ca^2+^]) from 9.0 to 4.0. Tissues were allowed to reach steady-state force (F) at each pCa. F-pCa curves were collected and analyzed with custom code using LabView software and were fitted to the Hill equation to calculate pCa_50_, the pCa at half-maximal force, and the Hill coefficient (n_H_), a measure of the cooperativity of force (30). The rate of tension redevelopment (*k*_tr_) was measured at maximal pCa and calculated from the half time of force recovery following a 15% rapid release−restretch transient. To measure tissue stiffness, high-frequency stiffness (HFS) was measured by applying a 1000 Hz sinusoidal length change of ± 0.5% of the muscle length (ML) and was calculated from the ratio of peaks of the Fourier transform of the force and ML signals.

To determine the effect of inorganic phosphate (P_i_) inhibition on demembranated tissue, the same F and *k*_tr_ measurements were collected as described above, in the same physiological solutions at pCa 9.0 and pCa 4.5 (21 °C) with incorporation of either 0, 1, 3, or 10 mM [P_i_] (30).

### Myofibril mechanics

Slices of minipig cardiac tissue were permeabilized overnight at 4 °C in rigor solution containing 2 mM DTT and 1% (v/v) Triton X-100. The following day, demembranated tissues were rinsed twice in rigor solution (2 mM DTT, 1:100 dilution of protease inhibitor (Sigma)) before being homogenized for 10 s at low speed with a Tissue Ruptor II and then stored on ice until use. Myofibril activation and relaxation measurements were performed on a custom-built setup as previously described (31, 32). Briefly, myofibrils were mounted between two glass needles, one a force transducer with a known stiffness (7.7 nN/μm) and the other an inflexible motor arm. A dual diode system measured force based on needle deflection. A double-barreled glass pipette delivered relaxing (pCa = 9.0) and maximal activating (pCa = 4.5) solutions to the mounted myofibril. To investigate changes in ADP release from myosin during the slow phase of relaxation, myofibrils were activated and relaxed in pCa 4.5 and pCa 8.0, respectively, and then paired measurements were performed in pCa 4.5 and pCa 8.0, with 50% of the 5 mM nucleotide pool replaced with ADP (50:50 ADP:ATP, each at 2.5 mM). Activation and relaxation data were collected at 21 °C and analyzed as previously described (32).

### Molecular-dynamics simulations

*Model Preparation.* The model of the post-powerstroke-like actomyosin cross-bridge was based on multiple cryo-EM structures solved by Doran *et al*. (PDB ID: 8EFH) (33). The authors of the cryo-EM structure presented three structures in total: (1) a particle-averaged structure of rigor-like myosin + ELC (PDB: 8EFD), (2) a helically reconstructed structure of post-powerstroke myosin (myosin.ADP.Mg^2+^) in complex with a thin filament (actin 5mer + tropomyosin) (PDB: 8EFH), and (3) a single-particle reconstruction of post-powerstroke myosin.ADP.Mg^2+^ + ELC complex (PDB: 8EFE). In the helically averaged structure, the tail and ELC were not modeled. This was also the case in the authors’ model of the rigor-state complex. Our model is based on these structures. We retained the F-actin pentamer, and the tropomyosin strands were removed. The missing N-terminal residues for each actin monomer (CDDEE) were built using *Modeller* (34). We created hybrid structures for myosin by combining myosin chains from 8EFH and 8EFE. Residues 6 – 702 from the 8EFH structure were retained for the motor domain (to define appropriate interactions with the thin filament), and residues 703 – 796 from 8EFE were grafted onto it. Coordinates for missing residues were built using *Modeller*. The ELC was modeled based on its structure in 8EFE. Residues 707 – 850 of myosin and the RLC residues were built via homology modeling using PDB: 3I5G (35) as a template and grafted onto our system. Our final model includes a thin filament pentamer, β-myosin.ADP.Mg^2+^, the ELC, and the RLC. His protonation states at pH 7.0 were predicted using the H++ webserver (36).

*Molecular Simulation.* The resulting system was prepared for molecular dynamics simulation using the AMBER20 simulation package (37). Hydrogen atoms were modeled onto the initial structure using the leap module of AMBER, and each protein was solvated with explicit water molecules in a truncated octahedral box, and ∼120 mM KCl counterions were added. The hydrogen mass repartitioning scheme was employed to enable a 4-fs timestep (38). Protein atoms were modeled with the ff14SB force field, water molecules were modeled with the TIP3P force field (39), metal ions were modeled using the Li and Merz parameter set (40–42), and ADP molecules were treated with the GAFF2 force field (43). Partial charges for ADP were derived from a restrained electrostatic potential fit to quantum mechanics calculations performed using ORCA (44). Each system was minimized in three stages. First, hydrogen atoms were minimized for 1,000 steps in the presence of 25 kcal/mol restraints on all heavy atoms. Second, all solvent atoms were minimized for 1,000 steps in the presence of 25 kcal/mol restraints on all protein atoms. Third, all atoms were minimized for 8,000 steps in the presence of 25 kcal/mol restraints on all backbone heavy atoms (N, O, C, and Cα atoms). After minimization, systems were heated to 310 K using the NVT (constant number of particles, volume, and temperature) ensemble and in the presence of 25 kcal/mol restraints on backbone heavy atoms. Next, the systems were equilibrated over five successive stages using the NPT (constant number of particles, pressure, and temperature) ensemble. The systems were equilibrated for 5.4 ns in the presence of restraints on backbone heavy atoms that successively decreased from 25 kcal/mol to 0 kcal/mol. A 10 Å nonbonded cutoff was used for all preparation and production simulations. The equilibrated systems were then simulated using conventional molecular dynamics protocols in the NVT ensemble in triplicate for 500 ns each (6 total simulations at 500 ns each = 3 µs total sampling), and coordinates were analyzed every 10 ps.

*Molecular Analyses.* Residue-residue contacts in the simulations, interatomic distances, and nucleotide-myosin interaction energies were calculated with *cpptraj* (45). Two residues were considered in contact with one another if at least one pair of heavy atoms was within 5 L. Protein images were generated with UCSF Chimera (46).

### Human induced pluripotent stem cell generation, maintenance, and cardiomyocyte differentiation

Clustered regularly interspaced short palindromic repeats and CRISPR-associated protein 9 (CRISPR/Cas9) was used to induce the *MYH7* c.1208G>A (R403Q) mutation in a well-established (WTC-11) human induced pluripotent stem cell (hiPSC) line (derived from skin fibroblasts, male). 1×10^6^ WTC-11 hiPSCs were electroporated with Cas9 (0.3 μM, Sigma) and gRNA (1.5 μM, Synthego, TCATTGCCCACTTTCACCCG) as an RNP complex, along with 2 μM of single stranded DNA (ssDNA, AGTCTGCCTACCTCATGGGGCTGAACTCAGCCGACCTGCTCAAGGGGCTGTGCCACCCTCAGGTGAAAGTGGGCAATGAGTACGTCACCAAGGGGCAGAATGTCCAGCAGGTGGGTCCATC, IDT) using Amaxa nucleofector (Human Stem Cell Kit 2) in the presence of Y-27632 ROCK inhibitor (10 μM, Selleck Chemicals), with or without HDR enhancer (IDT). Individual colonies were hand-picked and plated into 96 well plates. DNA was extracted using QuickExtract™ DNA Extraction Solution (Epicentre #QE09050) and nested PCR was performed using custom primers: F: GGCAGCTGTCAAGTCATGGA and R: AGATCTCGAAGCCAGCGATG. The PCR product was purified using EXO-SAP enzyme (ThermoFisher) and sent for Sanger sequencing analysis (Genewiz). One representative hiPSC clone (clone 99), which was unsuccessfully gene-edited and wildtype for the *MYH7* gene sequence, was selected as our control line for this study (Supplemental Figure S4). One hiPSC clone (clone 62), confirmed to have a heterozygous R403Q mutation in *MYH7,* was used as our representative R403Q hiPSC line (Supplemental Figure S4). Both clones were analyzed for potential off target mutations in *MYH6* gene using PCR primers *MYH6* F: AACAGTGCTAGACAGGTGCC and *MYH6* R: AGGAGCAAGCGAGTGATTGT, followed by Sanger sequencing confirming the absence of mutation in this *MYH6* region. Selected hiPSC clones were expanded, confirmed to be negative for mycoplasma, and banked in liquid nitrogen.

The genomic allele copy number and wildtype/heterozygous status of the clones were confirmed by ddPCR using the QX200 Manual Droplet Digital PCR System (Bio-Rad) as per the manufacturer’s protocol. A custom ddPCR Mutation Detection assay containing one PCR primer set and two probes (one for WT (HEX) and one for mutant detection (FAM)) was designed by BioRad (S Figure S4).

The protocol for hiPSC maintenance and differentiation into human induced pluripotent stem cell-derived cardiomyocytes (hiPSC-CMs) was based on our recent publication (47). Briefly, a small molecule, WNT signal modulation protocol was used to differentiate hiPSCs that had been cultured on 1X Matrigel (Corning) in mTeSR1 media (STEMCELL™ Tech). CHIR99021 (Cayman Chemicals) and WNT C-59 (Tocris) were used between day 0 – 2 and 2 – 4 of the cardiomyocyte differentiation process, respectively. Basal media used between day 0 – 6 was Roswell Park Memorial Institute (RPMI) media supplemented with 213 μg/mL ascorbic acid 2-phosphate (Sigma) and 500 μg/mL bovine serum albumin (BSA, Sigma). After day 6, basal media was changed to RPMI and 1X B27 supplement (ThermoScientific), denoted as cardiomyocyte maintenance media. On day 14, wells with >70% visually beating hiPSC-CMs were replated in replating media (RPMI + 1X B27 supplement + 0.5% FBS + 10 μM Y-27632 ROCK inhibitor) for lactate purification. Lactate purification (4 mM sodium L-lactate, Sigma) in glucose-free RPMI basal media (ThermoFisher) was initiated on day 15 and continued for four days. Starting on day 19, hiPSC-CMs were returned to cardiomyocyte maintenance media for long term culture.

### Cell patterning, myofibril isolation and myofibril mechanics

To perform hiPSC-CM myofibril kinetics experiments, hiPSC-CMs were cultured for 7 days (day 35 – 45 post-initiation of cardiomyocyte differentiation) on lined patterns of stamped Matrigel to elongate and mature hiPSC-CM myofibrils prior to myofibril isolation, as recently published (47). Briefly, PDMS stamps with the lined pattern geometry were coated with 5X Matrigel overnight, stamped onto an 18 mm coverglass, and subsequently placed stamp-side down onto a 10 kPa polyacrylamide gel crosslinked with 10% ammonium persulfate (APS, Sigma) and TEMED (BioRad) which had been pipetted onto a plasma-treated, bind-silane coated, glass-bottom 6-well culture dish. On ∼ day 38 post-initiation of differentiation, ∼ 5×10^5^ hiPSC-CMs were replated onto this patterned surface in modified replating media (RPMI + 1X B27 supplement, and 10 μM Y-27632 ROCK inhibitor). To elevate the basal media’s calcium concentration to 1.1 mM, patterned hiPSC-CMs were fed with RPMI and DMEM minus glucose in a 50:50 ratio with 1X B27 supplement. Media was changed every 48 hours for 1 week until the day of myofibrils isolation (hiPSC-CMs 7 days on patterned surface).

At day 42 – 48 post-initiation of differentiation, myofibrils were isolated from cells cultured on patterned surfaces by permeabilizing the tissue in pCa 8.0 relaxation solution (170 mM ionic strength, 3 M pMg, 15 mM total EGTA, 80 mM total MOPS, 5 mM MgATP, 15 mM total creatine phosphate, 83 mM free K, and 52 mM free Na, pH 7.1 at 4 °C) with 1:100 protease inhibitor cocktail (PIC, Sigma) and 1% (v/v) Triton X-100 (Acros Organics) for 10 mins on ice (47, 48). After washing twice with pCa 8.0 containing PIC, myofibrils were collected from the plate using a cell scraper and transferred into a tube. To separate bundles of myofibrils, the myofibril solution was gently homogenized ∼10 – 15 times using a Dounce tissue grinder pestle (Sigma).

Isolated myofibrils were then plated onto our custom-built myofibril apparatus, and individual myofibrils were picked up from the glass microscope surface between two glass microprobes (21 °C). One microprobe was controlled as a motor arm to set myofibril length to 2.3 μm, and the second microprobe acted as an optical-based force probe with a known stiffness (7.7 nM/nm). A double-barreled pipet enabled rapid Ca^2+^ switching, allowing measurement of myofibril force and contraction/relaxation kinetics at the millisecond timescale (47–49). To achieve maximal activation, all myofibrils were activated with pCa 4.5. To investigate changes in ADP release from myosin during the slow phase of relaxation, myofibrils were activated and relaxed in pCa 4.5 and pCa 8.0, respectively, with 50% of the 5 mM nucleotide pool replaced with ADP (50:50 ADP:ATP, each at 2.5 mM) (47).

### EHT generation, culture, and intact contraction experiment

Casting and maintenance of EHTs from hiPSC-CMs and HS27a human bone marrow stromal cells followed a protocol from Bremner *et al*. (50). Day 24 hiPSC-CMs from both cell lines and HS27a stromal cells were prepared in cardiomyocyte maintenance media (RPMI 1640 and 1X B27 Supplement). Appropriate volumes of cardiomyocytes and stromal cells were combined to achieve a cell count of 5×10^5^ hiPSC-CMs and 5×10^4^ HS27a cells per EHT. These cells were then resuspended in a volume of cardiomyocyte maintenance media sufficient to achieve 87 μL per EHT. The mold for the EHT was cast into 2% agarose, aliquoted into a 24-well dish, using a 3D-printed polylactic acid (PLA) spacer gifted by Dr. Nathan Sniadecki, University of Washington. The rack of PDMS posts with one flexible and one rigid post was also gifted to us by Dr. Sniadecki. This rack of posts was placed into the agarose EHT mold, and the cell mixture of 87 μL + 10 μL fibrinogen + 3 μL thrombin per EHT was pipetted into the mold to cast around each set of PDMS posts for 90 mins. The fibrin-gel encapsulated hiPSC-CMs and stromal cells, now adhered between the PDMS posts, were then lifted out of the agarose mold and transferred to a fresh 24-well plate with EHT culture media (RPMI 1640, 2% B27 Supplement, 5 g/L aminocaproic acid). Media was refreshed every 48 hours.

After three weeks in culture (day 45 post-initiation of hiPSC-CM differentiation), intact isometric contraction measurements were performed using the IonOptix™ Intact Muscle Chamber system. EHTs were cut off the PDMS posts and clipped between two, 1 mm wide platinum ribbon Omega clips. The EHT could then be suspended between length controller and force transducer probes within the IonOptix™ Intact Muscle Chamber system with circulating and oxygenated DMEM/F12 media supplemented with calcium to achieve 1.8 mM Ca^2+^. Media bath was maintained at 30 – 33 °C. EHTs were stretched to an optimal sarcomere length by stretching the tissue to maximize peak twitch amplitude. Intact twitch recordings were collected at 1 Hz pacing frequency (10 V stimulation and 10 ms duration). Twitch recordings were analyzed using the IonOptix IonWizard software. Force was normalized to EHT cross-sectional area assuming an elliptical shape.

### Myosin isoform stoichiometry

Protein lysates were isolated from both porcine myofibril samples (*N =* 3) and day 45 hiPSC-CMs (*N =* 2 – 3). For porcine myofibril samples, frozen pellets were thawed on ice and resuspended in sample lysis buffer (5% SDS, 0.75% sodium deoxycholate, 50 mM Tris-HCl pH 8.5, and 1:100 PIC). Samples were boiled at 95 °C for 10 mins, centrifuged at max speed for 15 mins, and the supernatant was collected and stored in a fresh tube prior to freezing at –80 °C. For hiPSC-CMs, cells cultured on a flat plastic substrate in basal cardiomyocyte media for 45 days were collected, spun down at 1,000 rpm for 5 min, and flash-frozen in liquid nitrogen before storage at –80 °C. At a later date, hiPSC-CMs cell pellets were thawed on ice, resuspended in 200 μL of the same sample lysis buffer, boiled at 95 °C for 10 mins, and centrifuged again at maximum speed for 15 mins. The resulting supernatant was stored at –80 °C for future experiments. Protein concentrations for both porcine and hiPSC-CM samples were determined using the Pierce™ BCA Protein Assay Kit (Thermo Scientific).

For myosin isoform analysis, gels were cast with a 6% acrylamide resolving gel (37.5:1 acrylamide:DATD) and a 2.95% acrylamide stacking gel (5.6:1 acrylamide:DATD) using mini gel plates with a 0.75 mm gel spacer and comb. TEMED and 10% APS were used as crosslinking agents. Protein lysates containing 0.55 μg of protein for porcine samples or 10 μg for hiPSC-CMs were mixed 1:1 with 2X Laemmli buffer (BioRad) containing β-mercaptoethanol (β-ME), boiled at 95 °C for 30 s, vortexed, and boiled again at 95 °C for 10 mins. Running buffer was composed of 50 mM Tris base, 0.384 M Glycine, and 0.1% (w/v) sodium dodecyl sulfate (SDS). Running buffer contained within the gel running cassette contained 0.14% β-ME. Gels were run for ∼2 hours at 4 °C, at 16 mA per gel, with constant stirring. Gel electrophoresis was stopped manually when the 150 kDa ladder band (BioRad Precision Plus Dual Color Protein Ladder) disappeared from the gel. Gels were then removed from the casting plates and fixed for 1 hour in 45% methanol and 10% acetic acid at room temperature with constant agitation. Myosin bands were visualized by overnight staining with 1X One-Step Lumitein™ Protein Gel Stain (Biotium) at room temperature with constant agitation, protected from light. The next day, gels were destained twice in ultrapure water for 5 mins, and imaged using a BioRad ChemiDoc under SYPRO Ruby protein stain settings.

### Phosphorylation analysis

Protein lysates isolated from both porcine and hiPSC-CM samples were also used to quantify phosphorylation of key sarcomeric proteins of interest (cMyBP-C, cTnI, and RLC). 10 μg of protein from each sample were combined 1:1 with 2X Laemmli buffer containing β-ME, boiled for 5 mins at 95 °C, and spun down at maximum speed for 5 mins. Samples were then loaded onto a 4 – 20% gradient MiniPROTEAN® TGX Stain-Free Protein Gel (BioRad). Tris-glycine running buffer was prepared with 0.1% (w/v) SDS. The BioRad Precision Plus Dual Color Protein Ladder was used at a 1:25 dilution in 2X Laemmli buffer. Gels were run at 250 V until the dye front reached the bottom. Phosphorylation was quantified using the Pro-Q™ Diamond Phosphoprotein Gel Stain Kit (Invitrogen) following the manufacturer’s protocol. Briefly, gels were fixed in 50% methanol and 10% acetic acid twice for 30 mins each, washed in ultrapure water three times for 10 mins, and then stained with Pro-Q™ Diamond stain for 90 mins at room temperature with constant agitation while protected from light. Stained gels were then destained (20% acetonitrile, 50 mM sodium acetate, pH 4.0) three times for 30 mins and washed in ultrapure water twice for 5 mins. Gels were imaged using a BioRad ChemiDoc with the Pro-Q™ Diamond settings. To quantify total protein, gels were subsequently stained overnight with SYPRO® Ruby Protein Gel Stain (Invitrogen) and then destained for 30 mins (10% methanol, 7% acetic acid). SYPRO® Ruby-stained gels were imaged using the SYPRO® Ruby settings on the BioRad ChemiDoc. Gel analysis was performed using ImageJ2 (version 2.14.0/1.54j). Densitometry of Pro-Q™ Diamond phosphorylation bands was normalized to corresponding total protein bands from the SYPRO® Ruby stain. Porcine samples were analyzed for cMyBP-C, cTnI, and RLC phosphorylation. HiPSC-CM samples were analyzed for cMyBP-C and RLC phosphorylation, but not cTnI, as no cTnI was detected at day 45 in a previous HCM hiPSC-CM line studied in our lab (47).

### Statistics

Data are presented as mean ± standard error of the mean (SEM). Statistical analyses comparing technical replicates within each group were performed using GraphPad Prism (version 9.4.1). A threshold for statistical significance was set at *p <* 0.05. Normality was assessed using the Shapiro-Wilk test. For data that were normally distributed, statistical comparisons were made using either an unpaired, two-tailed Student’s t-test with Welch’s correction or a two-way ANOVA with Tukey’s multiple comparisons test. For non-normally distributed data, the nonparametric Mann-Whitney test was used.

## IV. RESULTS

### Effect of the R403Q mutation on myosin populations in relaxed porcine ventricular muscle

Myosin heads that are free from the thick filament backbone have higher ATPase activity and make up the readily available pool of myosins that can contribute to cross-bridge formation and cycling during contraction. We used two approaches to determine how this population is affected by the R403Q mutation: low-angle X-ray diffraction and super-resolution single-molecule fluorescence microscopy.

***X-Ray diffraction*.** In the equatorial plane, the ratio of the intensities of the 1,1 and 1,0 reflections, I_1,1_/I_1,0_, reports the radial position of myosin heads relative to actin filaments in sarcomeres (17, 51). Example X-ray patterns for WT, R403Q, and R403Q with 2-deoxy-ATP (dATP: a strong myosin activator) (18, 52) in place of ATP in resting myocardium (pCa 8.0) are shown in Figure 1A and Supplemental Figure S1A−C. There was no significant difference in I_1,1_/I_1,0_ between WT (0.28 ± 0.01) and R403Q (0.27 ± 0.02), but the ratio was significantly increased in R403Q with dATP treatment (0.34 ± 0.02). Inter-filament lattice spacing (d_1,0_) was significantly greater in R403Q (38.83 ± 0.28 nm) than in WT (37.99 ± 0.22 nm) and was not further increased by dATP (Figure 1C). In the meridional plane, the intensities of the myosin-based layer line (I_MLL1_) and third-order myosin reflection (I_M3_) reflect the degree of ordering of myosin heads on the thick filament backbone. I_MLL1_ and I_M3_ were significantly lower in R403Q myocardium for both ATP and dATP conditions (Figure 1D,E). The spacing of the sixth-order myosin based reflection (S_M6_) increases when myosin heads shift from the OFF to ON state (17, 18, 53). S_M6_ was significantly larger in R403Q myocardium (7.261 ± 0.001 nm) than in WT myocardium (7.243 ± 0.002 nm) (Figure 1F). Replacement of ATP with dATP did not result in a further change. Taken together, the data suggest that the R403Q mutation reduced the ordering of myosin heads on the thick filament, facilitating the OFF-to-ON transition without significantly effecting the intensity ratio. This was accompanied by increased inter-filament lattice spacing, without myosin heads moving significantly away from the thick filament backbone toward the thin filaments.

**Figure 1.**
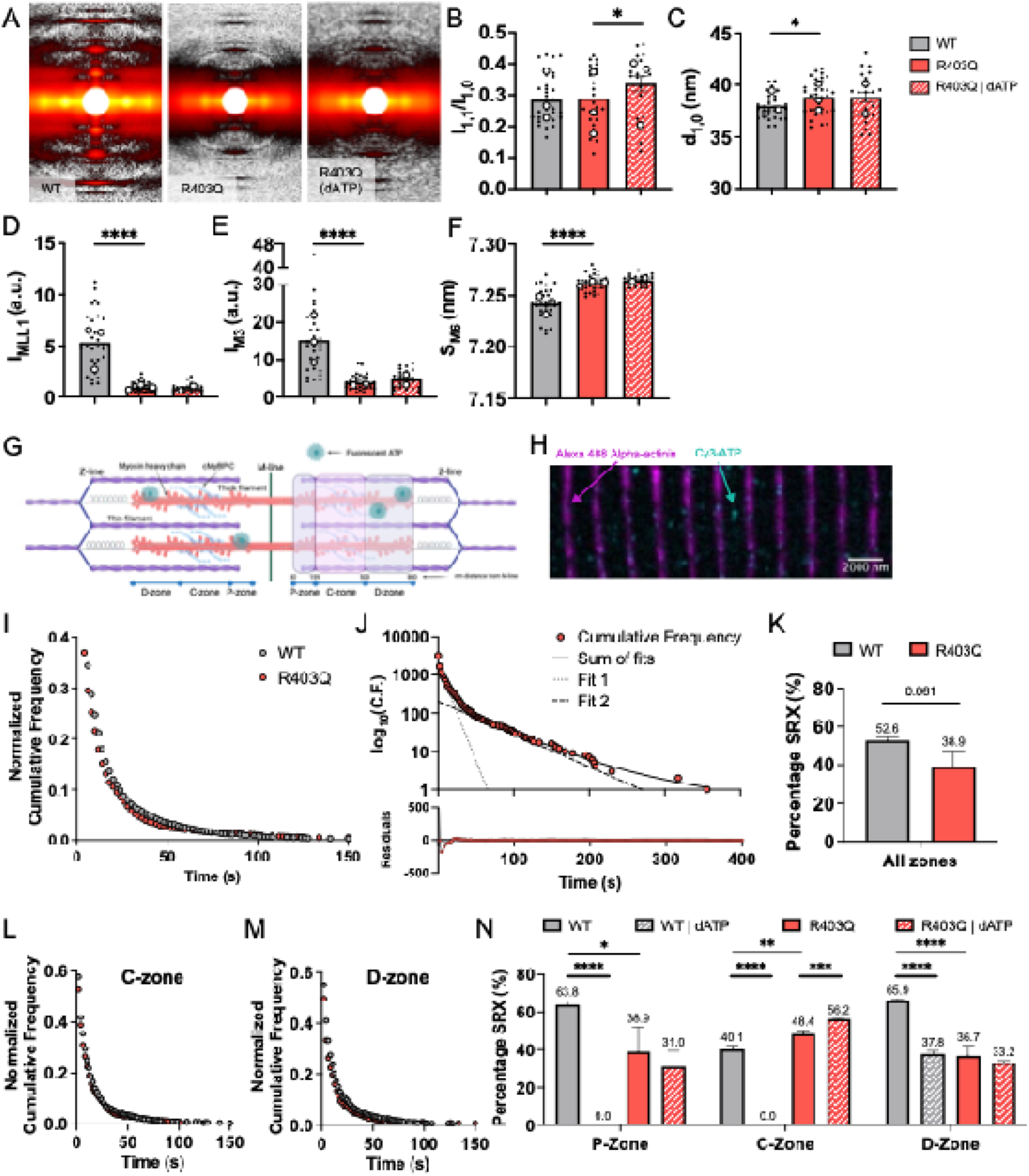
R403Q mutation elevated both structural and biochemical myosin recruitment in porcine myocardium. (A) Sample X-ray diffraction patterns (uncropped patterns in Supplemental Figure S1) for WT, R403Q, and R403Q tissue treated with 100% dATP nucleotide under relaxed conditions during small-angle X-ray exposure. Summary data for (B) the intensity ratio of the primary equatorial reflections (I_1,1_/I_1,0_), (C) lattice spacing (d_1,0_), (D) intensity of the first-order myosin-based layer line (I_MLL1_), (E) intensity of the third-order myosin-based meridional reflection (I_M3_), and (F) spacing of the sixth-order myosin-based meridional reflection (S_M6_). Paired measurements of R403Q porcine tissue were performed with and without 100% dATP in the relaxation solutions (WT: *N =* 3, *n =* 30 – 31; R403Q: *N =* 3, *n =* 26 – 35; R403Q | dATP: *N =* 2 – 3, *n =* 17 – 23). *N =* number of biological replicate minipigs; *n =* number of technical replicate X-ray diffraction patterns analyzed. (G) Schematic showing the cardiac myofibril sarcomere, illustrating key sarcomere proteins and nanometer spacing of the P-, C-, and D-zones of the thick filament relative to the M-line. (H) Representative R403Q porcine ventricular myofibril with overlay of Z-lines stained against α-actinin (magenta, 488 nm illumination) and fluorescently labeled Cy3-ATP molecules (cyan, 561 nm illumination) (scale bar = 2000 nm). (I) Normalized cumulative frequency graph of all Cy3-ATP binding lifetimes averaged across the entire sarcomere for both WT (grey circles) and R403Q (red circles) myofibrils. (J) Same cumulative frequency data for R403Q myofibrils with frequency transformed onto a logarithmic scale (top). Dashed lines show individual exponential fits for DRX (Fit 1) and SRX (Fit 2) populations, which make up a cumulative fit of three exponentials (Sum of fits, black solid line). Residuals for the sum of the three exponential fits are shown for the R403Q data (bottom). (K) Percentage of myosin heads quantified to be in the SRX biochemical state in WT and R403Q myofibrils, averaged across the entire thick filament. Normalized cumulative frequency graphs of Cy3-ATP binding lifetimes specifically in the (L) C-zone and the (M) D-zone for both WT and R403Q myofibrils. (N) Percentage of myosin heads quantified to be in the SRX state, broken down by P-, C-, and D-zones of the thick filament for WT and R403Q myofibrils in the presence of either unlabeled 5 mM ATP or 5 mM dATP (both using 10 – 15 nM Cy3-ATP; WT: *N =* 3, WT | dATP: *N =* 3, R403Q: *N =* 3, R403Q | dATP, *N =* 2). The number of Cy3-ATP binding events for ‘all zones’ and for each sub-zone are listed in Supplemental Table S1. *N =* number of biological replicate minipigs. Data are shown as mean ± SEM. Significance calculated with Student’s t-test, \**p <* 0.05, \*\**p <* 0.01, \*\*\**p <* 0.001, \*\*\*\**p <* 0.0001.

***Single-molecule ATP tracking*.** To further investigate the impact of the R403Q mutation on myosin structure in resting muscle, we employed a single-molecule fluorescence approach using isolated cardiac myofibrils. Alpha-actinin stained Z-lines were used as the point of reference for the edges of sarcomeres (Figure 1G,H). Only myofibrils with an average sarcomere length of ≥1.7 μm were used for analysis to ensure that the integrity of the sub-zones of the sarcomere was not compromised (Supplemental Figure S2). The exact location with respect to the fluorescent Z-lines and duration of binding of fluorescently labeled Cy3-ATP molecules were determined (Figure 1H). An overlay of the normalized cumulative frequency graph for all events across the entire sarcomere is presented in Figure 1I. ATP binding lifetime data across the entire sarcomere and in each sub-sarcomere zone were fitted to the sum of three exponentials by minimizing the natural log of the sum square difference. Error analysis of the fit residuals (Figure 1J) indicated that three processes produced the optimal fit for both WT and R403Q datasets. Taking the natural log of the cumulative distribution data best illustrates the underlying exponential fits. The fastest population was not analyzed further since it was consistent with non-specific binding of Cy3-ATP (Figure 1J) (24, 54, 55). The slower two rates were attributed to the DRX and SRX populations.

The data averaged across the entire sarcomere thick filament suggested a total decrease in the percentage of heads in a low ATPase state for R403Q myofibrils relative to WT myofibrils, though the difference was not statistically significant (*p =* 0.061) (Figure 1K, Supplemental Table S1). When the results were broken down to the sub-sarcomere zones, the reduction in low ATPase heads in the R403Q vs WT myofibrils was attributed specifically to the P-(38.9 ± 13.17% vs. 63.8 ± 2.08%, *p =* 0.017) and D-zones (36.7 ± 5.43% vs. 65.9 ± 0.93%, *p <* 0.0001) (Figure 1L−N, Supplemental Table S1). The C-zone, which contains cMyBP-C, showed a significant increase in low ATPase myosin heads (48.4 ± 1.97% vs. 40.1 ± 2.12%, *p =* 0.007) for R403Q vs WT myofibrils (Figure 1L-N, Supplemental Table S1).

To determine whether C-zone myosin was still regulated by cMyBP-C, we replaced the entire pool of unlabeled ATP in the imaging buffer with the strong myosin activator, dATP, while Cy3-ATP was maintained as the fluorescent ATP indicator. In WT myofibrils, dATP completely eliminated low ATPase myosin in the P– and C-zones and significantly reduced low ATPase myosin in the D-zone (Figure 1N). In contrast, R403Q myofibrils with dATP exhibited no further decrease in low ATPase myosin in the P-zone and the D-zone, but a slight increase in low ATPase myosin in the C-zone (Figure 1N). Subsequently, the impact of dATP across all zones was not significant for R403Q myofibrils (*p =* 0.207, Supplemental Table S1). Thus, single-molecule analysis revealed that the R403Q mutation reduced low ATPase myosin heads in regions not regulated by cMyBP-C, while dATP can eliminate low ATPase myosins in the cMyBP-C zone to increase the amount of myosin in higher ATPase states. Combined with the X-ray data, these results suggest that for sarcomeres with the R403Q mutation, the net increase in ON (higher ATPase) myosin-in P– and D-zones, countered by the moderate decrease of ON myosin in the C-zone, may explain why the intensity ratio I_1,1_/I_1,0_ (average of all myosin heads) is not increased for R403Q muscle in X-ray diffraction.

### Effect of the **β**-myosin R403Q mutation on contractile and cross-bridge properties in porcine ventricular muscle

***Demembranated tissue mechanics*.** To determine the relationship between increased myosin availability (DRX cross-bridges) in resting cardiac muscle and force developed during Ca^2+^ activation, we measured steady-state force and the rate of force redevelopment following a rapid release−re-stretch protocol with varying concentrations of Ca^2+^. Normalized force−pCa curves are shown in Figure 2A. Maximal specific force (pCa 4.5) was not significantly different between R403Q and WT controls (Figure 2B, Supplemental Figure S2), but the Ca^2+^ sensitivity of force (pCa_50_) was significantly greater in the R403Q tissue (Figure 2C), with no significant difference in the Hill coefficient (n_H_; Figure 2D). High-frequency stiffness analysis suggested no difference in the number of cycling cross-bridges at a given force level, indicating that increases in force are likely due to increases in the cross-bridge recruitment (Supplemental Figure S3). The enhanced sub-maximal force with the R403Q mutation suggests that at least some of the increased number of myosins made available in resting muscle can be recruited for force development. The rate of tension redevelopment, *k*_tr_, at maximal Ca^2+^ was slightly, but not significantly, reduced (Figure 2E), suggesting maximal cross-bridge cycling rate may be somewhat affected by the R403Q mutation.

**Figure 2.**
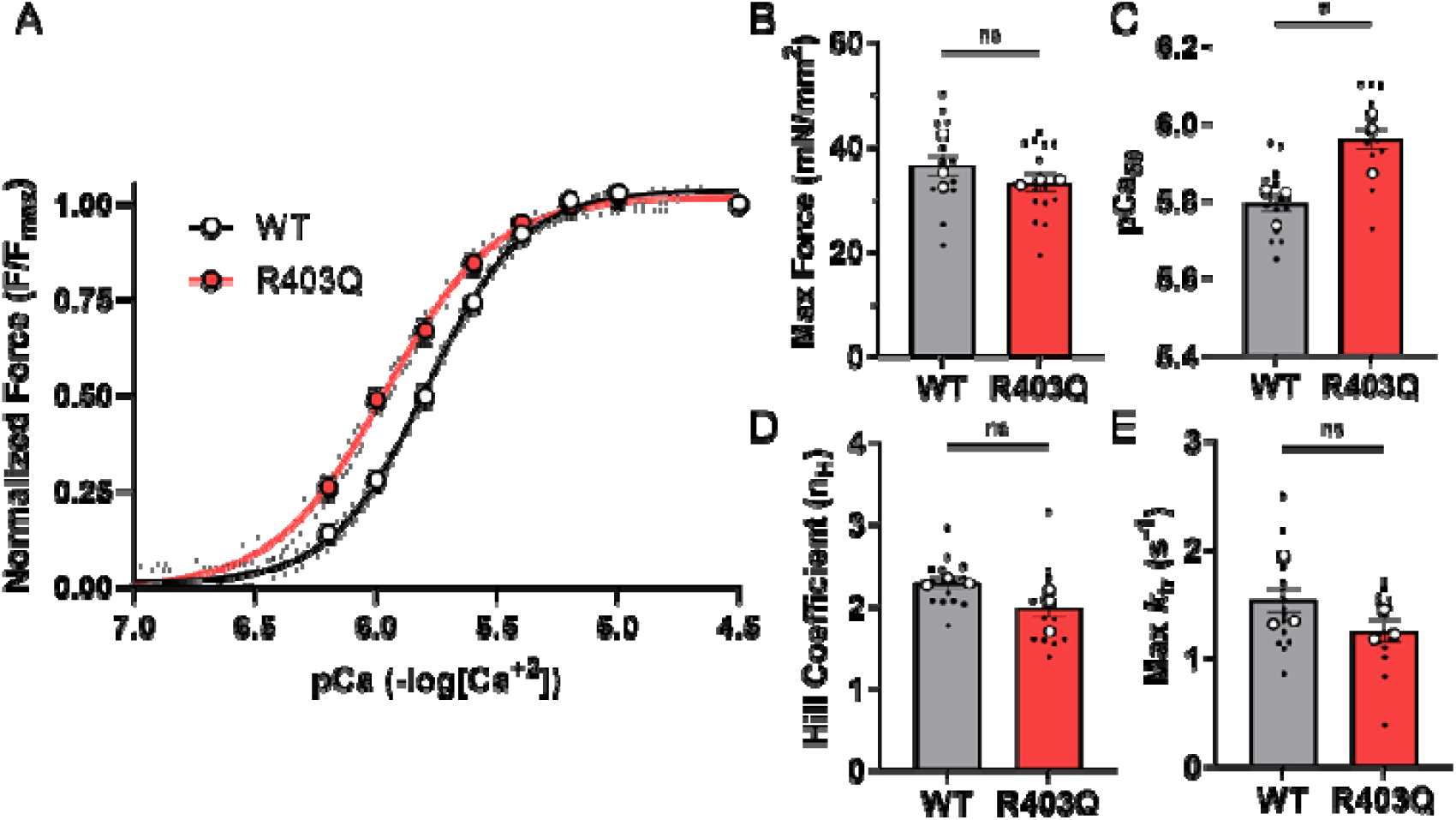
Isometric contraction of demembranated R403Q tissue reveals elevated calcium sensitivity. (A) Normalized force-pCa curves from permeabilized WT (black line) and R403Q (red line) porcine ventricular strips, and corresponding parameters of (B) force at maximal calcium activation solution (pCa 4.5), (C) pCa required for 50% maximum force (pCa_50_), (D) calculated Hill coefficient (n_H_), and (E) rate of tension redevelopment at maximal calcium (max *k*_tr_) (WT: *N =* 3, *n =* 14; R403Q: *N =* 3, *n =* 14). Data are presented as mean ± SEM. *N =* number of biological replicate minipigs; *n =* number of technical replicates. Unpaired, two-tailed Student’s t-test with Welch’s correction, \**p <* 0.05.

***Myofibril mechanics.*** To determine how the R403Q mutation impacts cross-bridge chemo-mechanics, we measured the kinetics of contraction and relaxation of single contractile organelles, i.e., myofibrils.

Isolated myofibrils were suspended between an optically based force transducer and a length-controlling lever arm at a sarcomere length of 2.3 μm. Representative force traces during the activation−relaxation protocol, switching from pCa 8.0 to 4.5, then back to 8.0, are shown in Figure 3A for both WT and R403Q myofibrils.

**Figure 3.**
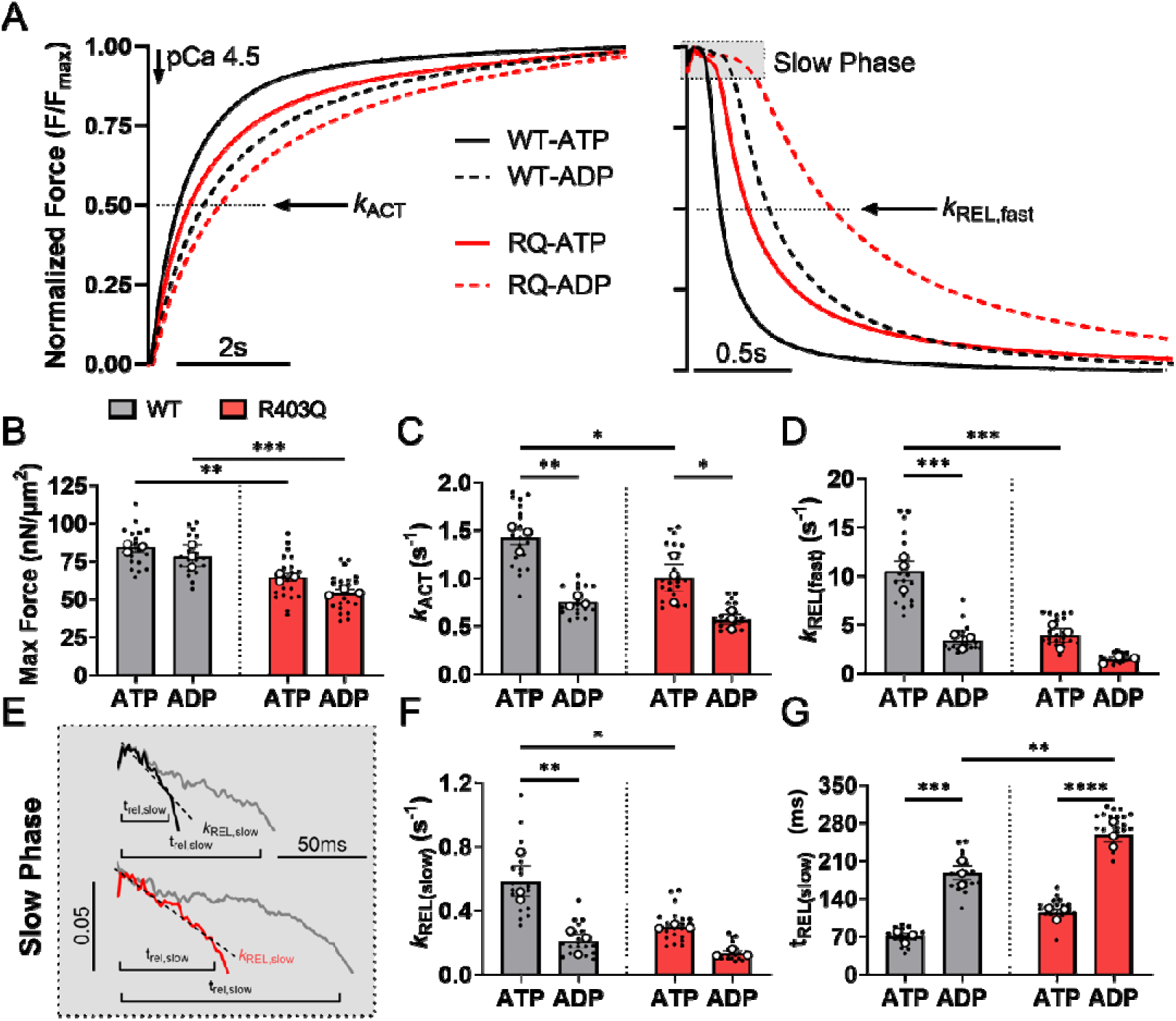
Porcine myofibrils with the heterozygous R403Q mutation exhibit reduced myofibril maximum force and slower contraction and relaxation kinetics. (A) Average normalized myofibril kinetics traces highlighting the exponential phases of contraction and relaxation for both WT (black line) and R403Q (red line) hiPSC-CM myofibrils. Paired myofibril traces were collected in the presence of 100% ATP and subsequently in the presence of 50% ATP and 50% ADP. Dashed lines indicate average normalized kinetics traces in the presence of elevated ADP. Contractile dynamics parameters at maximal pCa 4.5 include: (B) maximal force, (C) rate of activation (*k*_ACT_), (D) rate of the fast phase of relaxation (*k*_REL,fast_), (E) WT and R403Q traces (grey) highlighting the slow phase of relaxation (with a manual offset between the two cell lines for visualization) before and after the ADP inhibition experiment, (F) rate of the slow phase of relaxation under isometric length conditions (*k*_REL,slow_), and (G) duration of the slow phase of relaxation (t_REL,slow_) (WT: *N =* 3, *n =* 19; R403Q: *N =* 3, *n =* 22). Data are presented as mean ± SEM. *N =* number of biological replicate minipigs; *n =* number of myofibril technical replicates. Two-way ANOVA with Tukey post-hoc test, \**p <* 0.05, \*\**p <* 0.01, \*\*\**p <* 0.001, \*\*\*\**p <* 0.0001.

Maximal force (pCa 4.5) was significantly lower in R403Q myofibrils relative to WT (Figure 3B, Supplemental Table S2), and the rate constant (*k*_ACT_) of force development was slower (Figure 3C, Supplemental Table S2). The rate constant of the main, fast phase of relaxation (*k*_REL,fast_) was also significantly slower for R403Q myofibrils (Figure 3D, Supplemental Table S2). The initial phase of relaxation can be quantified based on two parameters: the rate constant derived from the slope of the linear phase (*k*_REL,slow_) and the duration of the linear phase (t_REL,slow_) (Figure 3E). *k*_REL,slow_ reflects the rate of cross-bridge detachment (56, 57), and t_REL,slow_ reflects the time it takes for thin filaments to deactivate (58). The *k*_REL,slow_ was slower for R403Q vs WT myofibrils (Figure 3F, Supplemental Table S2), suggesting the mutation slows myosin detachment kinetics, but t_REL,slow_ was not significantly different (*p =* 0.0658, Figure 3G, Supplemental Table S2).

Slower force development (*k*_ACT_) and relaxation (*k*_REL,slow_, *k*_REL,fast_) with the R403Q mutation could result from slower ADP release from myosin. ADP release can be probed with a product inhibition experiment, which was done by using activation and relaxation solutions containing a 50:50 ATP:ADP mixture (5 mM total). As expected, this slowed *k*_ACT_, *k*_REL,slow_, and *k*_REL,fast_, and prolonged t_REL,slow_ in WT myofibrils (Figure 3C-G). Interestingly, while elevated ADP also prolonged *k*_ACT_ and t_REL,slow_ for R403Q myofibrils, relaxation (*k*_REL,slow_, *k*_REL,fast_) was not significantly prolonged. This suggests that the R403Q mutation may result in modest impairment in ADP release associated with cross-bridge detachment, such that elevated ADP has little additional effect.

### Human models

***hiPSC-CM myofibril contractile properties***. The slower contractile kinetics of myofibrils with the R403Q mutation in pig ventricular muscle conflict with the faster kinetics of contraction and relaxation observed in myofibrils from myectomy samples from patients with the R403Q mutation (6, 7). Thus, to determine if differences were due to species or other factors, and to study the early stages of disease initiation, we engineered the heterozygous β-MHC R403Q mutation into WTC-11 human induced pluripotent stem cells (hiPSCs) using CRISPR/Cas9 gene editing (Figure 4A, Supplemental Figure S4), as described in the Methods. R403Q and WT cells were differentiated into cardiomyocytes (hiPSC-CMs), then replated onto Matrigel with a stamped lined pattern and cultured in the presence of elevated Ca^2+^ for 1 week (day 38 – 45 post initiation of differentiation) to enhance cell maturation and myofibril elongation (Figure 4B) (47). Isolated myofibrils were then mounted onto the myofibril apparatus between a length controller and a force transducer, and were stretched to 2.3 μm sarcomere length with visible sarcomeres (Figure 4C,D).

**Figure 4.**
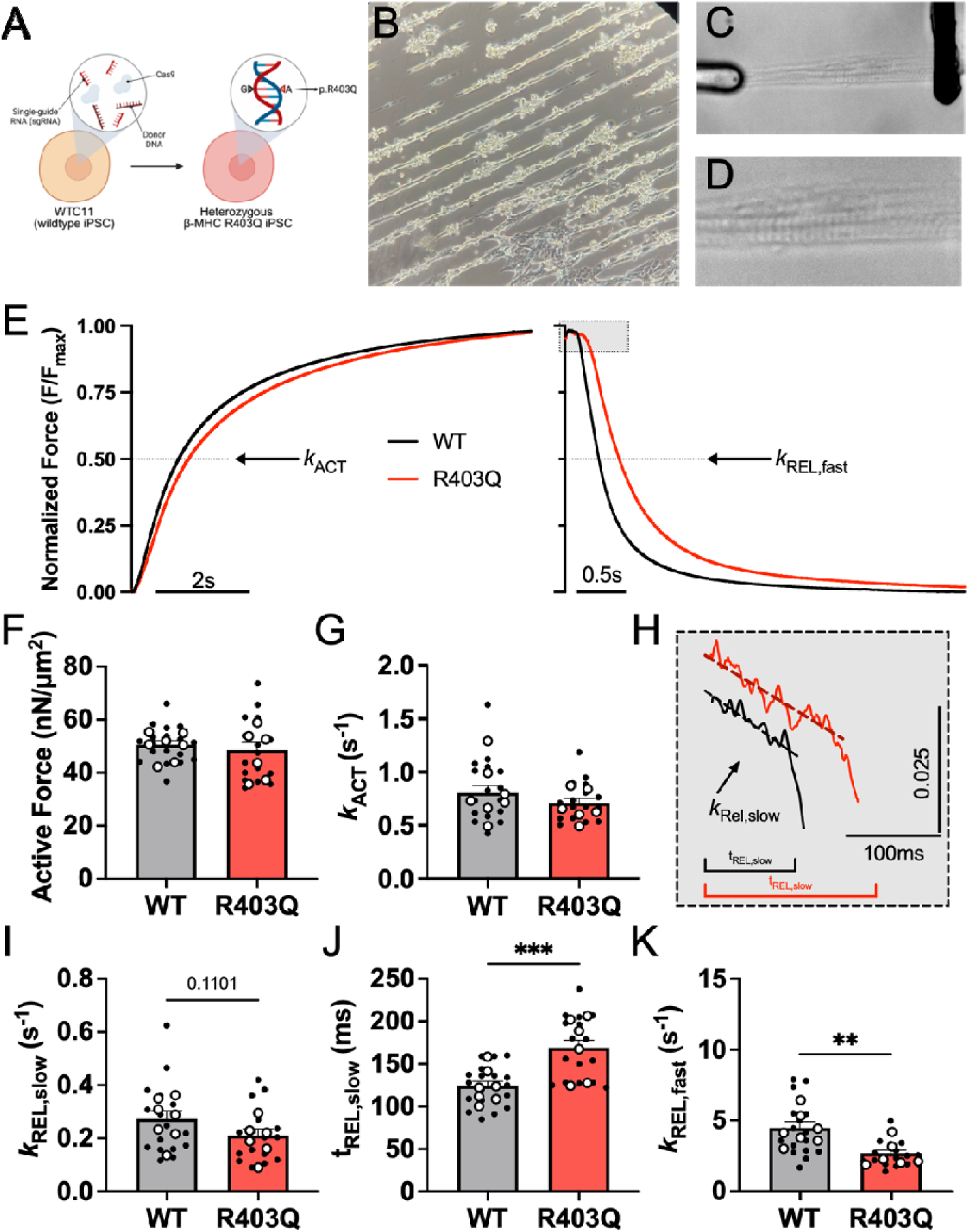
HiPSC-CM myofibrils with the heterozygous R403Q mutation exhibit impaired relaxation kinetics and prolonged thin filament deactivation. (A) Schematic illustration of CRISPR/Cas9 gene editing of WTC-11 hiPSCs to engineer the heterozygous R403Q mutation in the *MYH7* gene. (B) Representative image of hiPSC-CMs cultured on lined patterned Matrigel prior to myofibril isolation and (C) a bundle of isolated hiPSC-CM myofibrils mounted between two glass needles for mechanical measurements. (D) Magnified view of myofibril bundle highlighting sarcomere Z-lines. (E) Average normalized force traces showing the exponential phases of contraction and relaxation for both WT (black line) and R403Q (red line) hiPSC-CM myofibrils. Contractile dynamics parameters at maximal pCa 4.5 from day 42 – 48 hiPSC-CM myofibrils include: (F) maximal force, (G) rate of activation (*k*_ACT_), (H) WT and R403Q traces illustrating the slow phase of relaxation (manual offset for visualization), (I) rate of the slow phase of relaxation under isometric conditions (*k*_REL,slow_), (J) duration of the slow phase of relaxation (t_REL,slow_), and (K) rate of the fast phase of relaxation (*k*_REL,fast_) (WT: *N =* 7, *n =* 18 – 19; R403Q: *N =* 6, *n =* 16 – 17). Data are presented as mean ± SEM. *N =* number of biological replicate differentiation batches; *n =* number of myofibril technical replicates. Unpaired, two-tailed Student’s t-test with Welch’s correction, \*\**p <* 0.01, \*\*\**p <* 0.001.

Figure 4E shows normalized example force traces from both WT and R403Q hiPSC-CM myofibrils when maximally Ca^2+^ activated (pCa 4.5), then relaxed (pCa 8.0). The data for all measurements demonstrate that force was similar (Figure 4F, Supplemental Table S2), and *k*_ACT_ was not significantly different (Figure 4G, Supplemental Table S2) between WT and R403Q hiPSC-CM myofibrils. This differs somewhat from the porcine myofibril data, where both parameters were significantly reduced by the R403Q mutation. However, the relaxation kinetics (Figure 4H) agree better with the porcine data, with the R403Q myofibrils having reduced *k*_REL,fast_, a non-significant reduction in *k*_REL,slow_, and prolonged t_REL,slow_ (Figure 4I-K, Supplemental Table S2).

***EHT contractile properties*.** To determine how the R403Q mutation affects intact muscle twitches, we cast the hiPSC-CMs into fibrin scaffold-based EHT constructs affixed to PDMS posts. At the time of casting, WT EHTs had 99.1% hiPSC-CM purity and R403Q EHTs had 97.8% hiPSC-CM purity (data not shown). EHTs beat spontaneously for three weeks following seeding. For measurements, EHTs were removed from the PDMS posts and mounted between a force transducer and a length controller to measure isometric tissue force and kinetics of contraction and relaxation (Figure 5A). EHT cross-sectional area (CSA) was used to normalize peak force (Figure 5B). At 1 Hz pacing frequency, EHTs with the heterozygous R403Q mutation exhibited 50% greater twitch force than WT EHTs (Figure 5A,C). During the contraction phase, R403Q EHTs reached 50% force faster than WT EHTs (Figure 5D). However, the time to peak force was not significantly different (Figure 5E). The 50% relaxation time was not significantly different between WT and R403Q EHTs, but the 90% relaxation time was significantly prolonged (Figure 5F,G). Overall, these results demonstrate the heterozygous R403Q EHTs produce greater peak force and have impaired relaxation compared to WT EHTs. We also calculated the relative tension time index (TTI) (29, 59) between WT and R403Q EHTs. TTI is determined from the area under the twitch curve. When normalized relative to WT EHTs, TTI for R403Q EHTs was a positive value, which is predictive of a HCM hypercontractile phenotype (Figure 5H) (59).

**Figure 5.**
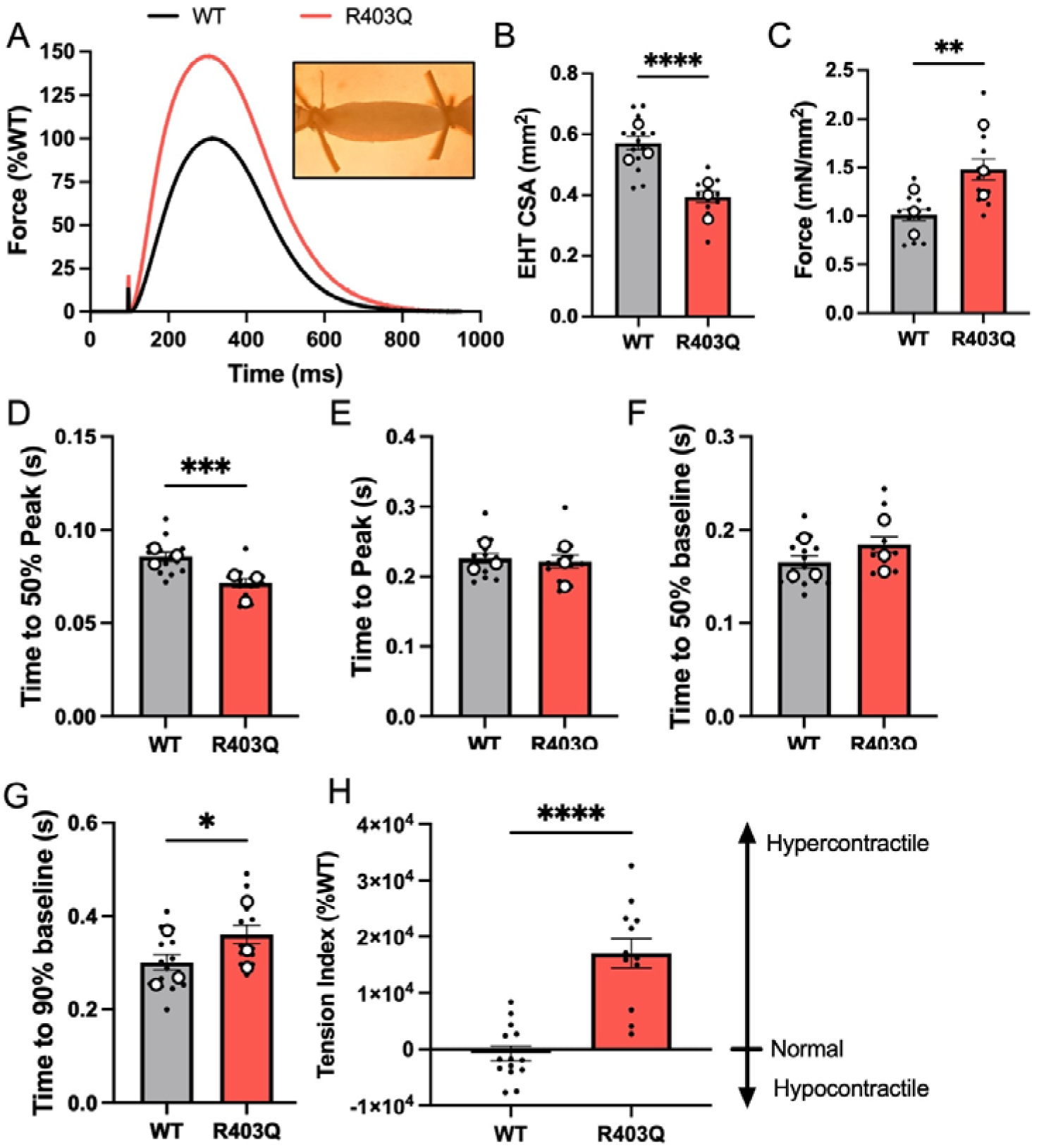
R403Q EHTs exhibit hypercontractility and impaired relaxation during isometric contraction. WT and R403Q hiPSC-CMs at day 24 post-initiation of differentiation were cast around one rigid and one flexible PDMS post in a 3D fibrin mold. EHTs matured and compacted around the PDMS posts for three weeks. At this three-week time point (day 45 post-initiation of differentiation), EHTs were removed from the PDMS posts and suspended between a force transducer and a length controller in an IonOptix™ Intact Muscle Chamber apparatus. (A) Average tissue force traces for WT (black line) and R403Q (red line) EHTs, with R403Q force values normalized relative to WT (set at 100%). Inset shows a representative image of an EHT mounted between the force transducer and length-controlling arms of the IonOptix™ system. (B) Average cross-sectional area (CSA) of EHTs and (C) twitch force normalized to CSA. (D) Time to 50% contraction, (E) time to peak force, (F) time to 50% relaxation, and (G) time to 90% relaxation. (H) Relative tension−time index (TTI), expressed as a percent of WT (set to zero) (WT: *N =* 3, *n =* 14; R403Q: *N =* 3, *n =* 12). All results were analyzed at a 1 Hz pacing frequency. Data are presented as mean ± SEM. *N =* number of biological replicate differentiation batches; *n =* number of EHT technical replicates. Unpaired, two-tailed Student’s t-test with Welch’s correction, \**p <* 0.05, \*\**p <* 0.01, \*\*\**p <* 0.001, \*\*\*\**p <* 0.0001.

### Molecular Dynamics (MD) simulations

To study changes in myosin structure that could result in altered interactions between actin and myosin that give rise to the observed changes in relaxation, we performed MD simulations of pre-powerstroke myosin and post-powerstroke actomyosin.

### Pre-powerstroke myosin

*Assessment of electrostatic surface potential*. MD simulations of pre-powerstroke myosin provide insight into the structure of myosin presented to actin (for binding) following ATP hydrolysis to ADP and inorganic phosphate. Simulations of pre-powerstroke human β-cardiac myosin (M.ADP.Pi) were performed for WT and R403Q. We used the NMRClust algorithm (60) as implemented in UCSF Chimera (46) to cluster the WT and R403Q simulations on residues involved in actin binding: the lower 50 kDa domain helix-loop-helix motif (residues 519−554), loop 4 (residues 363−376), and the cardiomyopathy loop (residues 401−415) (Figure 6A-C). NMRClust yielded a large number of lowly populated clusters with low intra-cluster RMSDs, indicating that the actin binding regions did not experience large-scale conformational changes in these simulations. The top three WT cluster representatives described 14%, 7%, and 6% of the total ensemble (Figure 6A); the top three R403Q cluster representatives described 7%, 6%, and 6% of the total ensemble (Figure 6B). Next, we compared the Cα RMSD of the top three cluster representatives for only the residues used to form clusters. The average RMSD was 1.6 Å among the WT representatives, 1.2 Å among the R403Q representatives, and 1.7 Å between the WT and R403Q representatives. Inspection of the representatives show that differences in WT/R403Q structure and clustering properties are attributable to differential sampling of the cardiomyopathy loop, as expected (Figure 6C). Next, we mapped the Coulombic electrostatic potential onto the surface for each WT (Figure 6D) and R403Q (Figure 6E) cluster representative. The R403Q mutation reduced positive electrostatic potential in the vicinity of the cardiomyopathy loop and also modulated the electrostatic potential of loop 4 but had minimal effects on the electrostatic potential of the helix-loop-helix motif. This suggests minimal impact on electrostatic interaction between pre-powerstroke myosin and thin filament actin. However, in the interacting-heads motif (IHM) state, R403 of the blocked head forms interactions with free head residues Y455 and Q454 (61) and in prior MD simulations, the blocked head cardiomyopathy loop formed numerous interactions with the HO-linker loop of the free head (62). Thus, altered electrostatic interactions due to R403Q may affect IHM stability more than actomyosin association.

**Figure 6.**
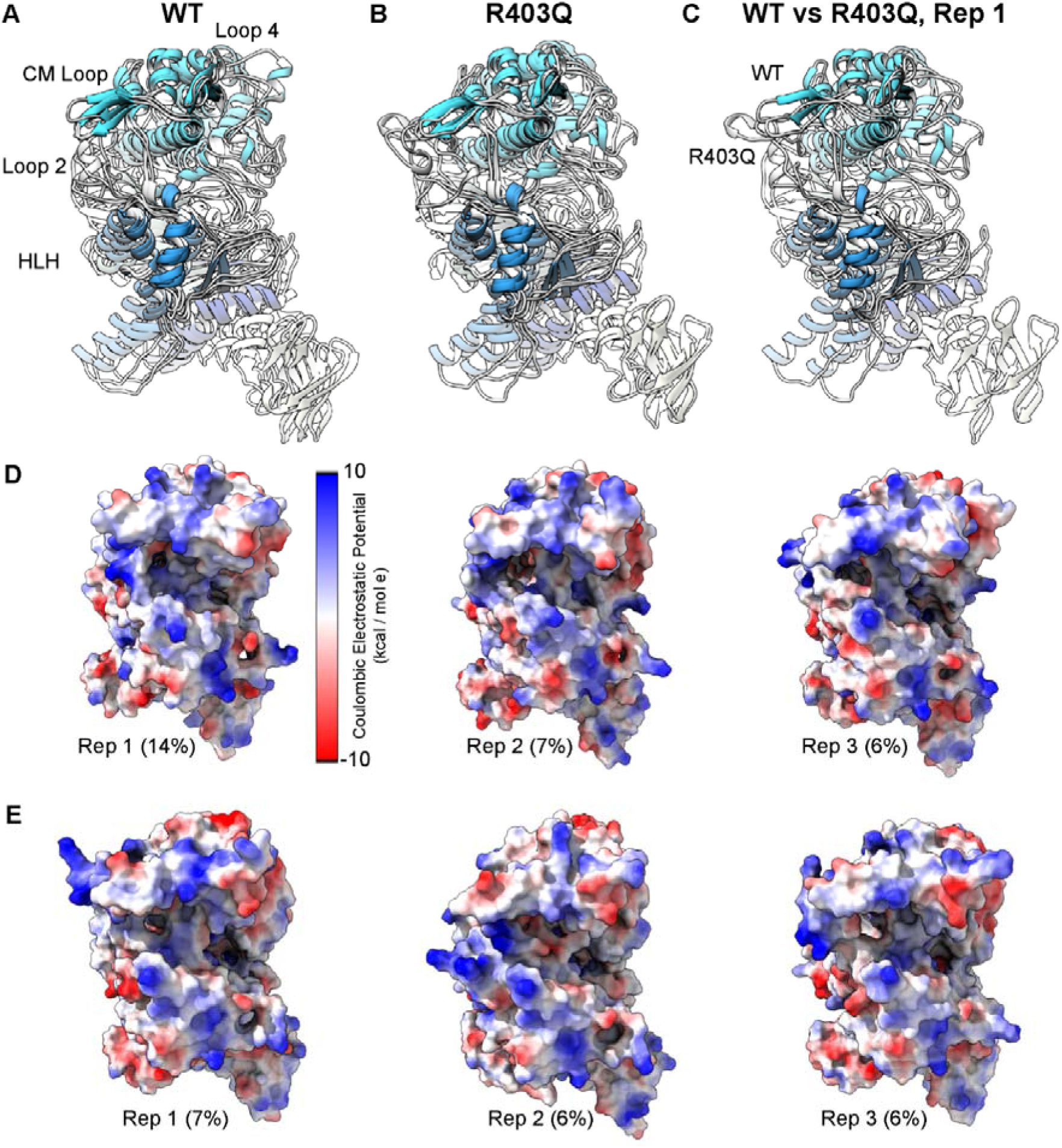
Effects of R403Q on the electrostatic surface potential of pre-powerstroke myosin. We extracted three representative myosin conformations from WT and R403Q pre-powerstroke myosin simulations. The (A) WT and (B) R403Q cluster representatives were then aligned to residues in the helix-loop-helix (HLH) motif, loop 4, and the cardiomyopathy loop (CM-loop). A comparison of the first WT and R403Q cluster representative (C) highlights that the major conformational difference is in the CM-loop. Mapping of the electrostatic potential onto the protein surface displays regions of net positive potential in blue and negative potential in red (D,E). Comparison of the WT (D) and R403Q (E) maps shows a decrease in positive potential in the CM-loop of the R403Q simulations.

*Post-powerstroke myosin-actin*. The slower relaxation of force in myofibrils with the R403Q mutation (assessed with *k*_rel,fast_, *k*_rel,slow_, and t_rel,slow_) and the reduced sensitivity of relaxation to elevated ADP suggest potentially altered post-powerstroke myosin-actin interactions. At the molecular scale, impaired relaxation may be attributed to altered ADP release, ATP binding, and/or the detachment of myosin heads from the thin filament. MD simulations of the post-powerstroke actomyosin complex (Figure 7A) investigated mechanisms by which the R403Q mutation impacts myosin-actin interactions. R403 is in the cardiomyopathy loop (CM-loop, residues 397 – 416), a β-hairpin in the upper 50 kDa domain of myosin that interacts with actin filaments when the myosin cleft is closed. In the cryo-EM structure, the CM-loop twists such that one side faces the thin filament and the other faces the myosin upper 50 kDa domain. We analyzed interactions made by the CM-loop and interactions at the actomyosin interface. The cryo-EM structure and the WT simulations indicate that R403 forms a salt bridge with E603 and that the aliphatic portion of the side chain stacks against Y410 (Figure 7B). R403Q modulated contacts between residues within the CM-loop (Figure 7C,D) as well as between myosin and actin. The Q403 – E603 electrostatic interactions were weaker, and the side chain did not stack as well against Y410 (Figure 7C). We also observed statistically significant differences in contacts made by the upper 50 kDa domain and loop 2 of myosin with the negatively charged N-terminus and the D24 – E31 loop of actin. Changes in residue-residue interactions oriented the CM-loop closer to the thin filament: for the WT simulations, the average distance between the CM-loop and the center of mass of the central actin monomer was 18.8 ± 0.3 Å; for the R403Q simulations, the average distance was 18.0 ± 0.1 Å (*p =* 0.01). We calculated the linear interaction energy at the actin-myosin interface and between the cardiomyopathy loop and actin for the WT and R403Q simulations. The R403Q mutation resulted in a stronger interaction between the cardiomyopathy loop and actin relative to WT (WT: –133 ± 43.8 kcal/mol; R403Q: –143 ± 41.5 kcal/mol; an 8% increase in interaction energy, errors are standard deviations), though this difference was not statistically significant. R403Q also increased the interaction energy between actin and myosin (WT: –2122 ± 213 kcal/mol; R403Q: –2328 ± 195 kcal/mol), a difference that was also not statistically significant. Note that this calculation does not account for the contributions of solvent to the interaction energy, the entropy of solvation, and only considers the post-powerstroke (AM.ADP*) conformation.

**Figure 7.**
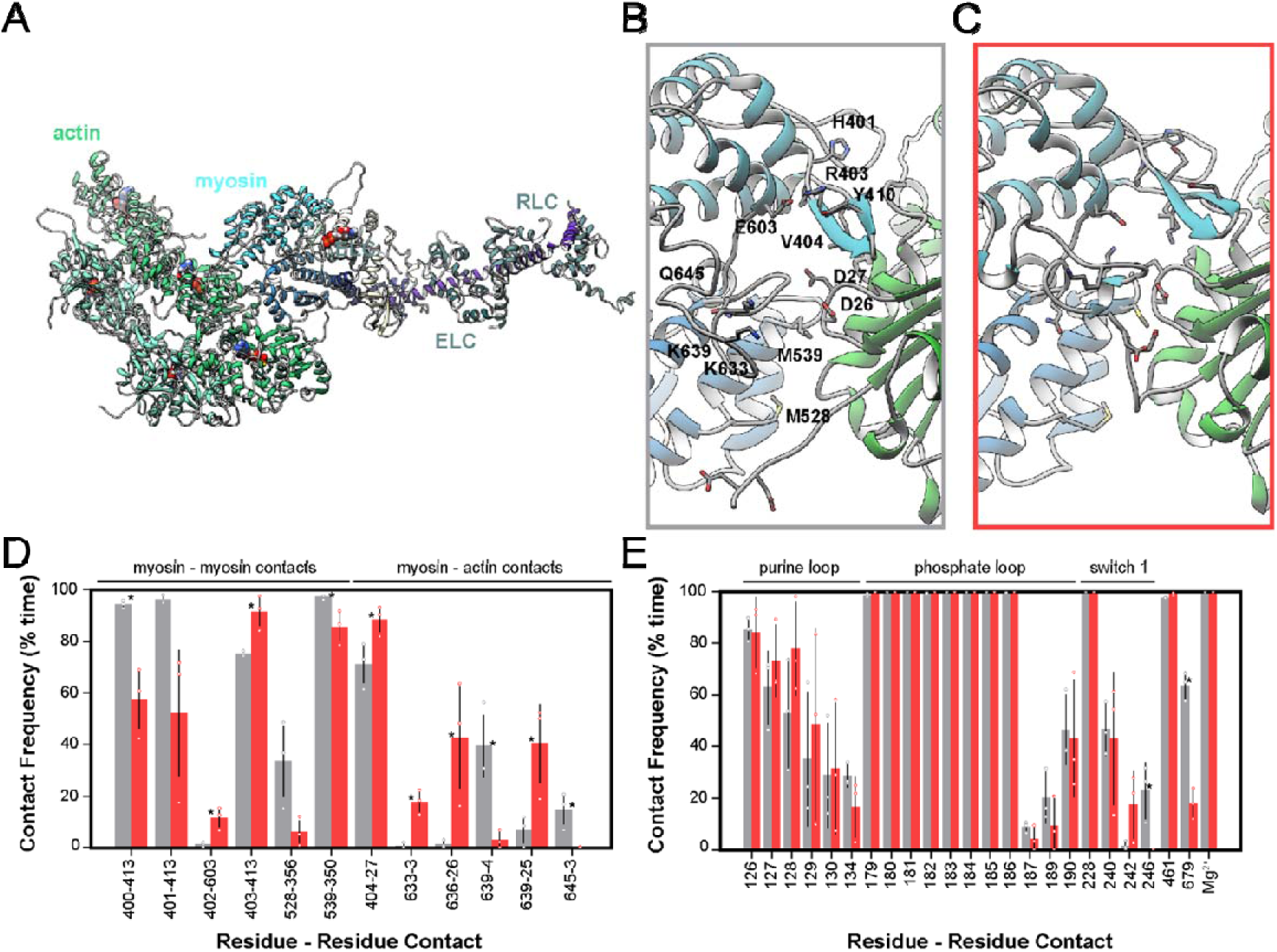
MD simulations of post-powerstroke WT and R403Q myosin. (A) Snapshot of the simulated system containing an actin pentamer, an ADP.Mg^2+^-bound myosin motor, the essential light chain (ELC), and regulatory light chain (RLC). Snapshots obtained from the (B) WT (grey) and (C) R403Q (red) simulations show that R403Q leads to altered interactions within the cardiomyopathy loop (CM-loop) and at the actomyosin interface. (D) Protein-protein contact frequencies (expressed as % simulation time) between myosin and actin residues in the WT (grey) and R403Q (red) simulations. (E) Contact frequencies (expressed as % simulation time) between myosin residues and ADP.Mg^2+^. Unpaired, two-tailed Student’s t-test, \**p <* 0.05.

Next, we analyzed interactions between the strongly bound myosin motors and ADP. Consistent with an ‘open’ conformation of the nucleotide binding pocket, there was large variation between simulations in interactions between ADP and the purine binding loop and switch 1 of myosin (Figure 7E). We observed few statistically significant changes in the interactions between myosin motors and their bound nucleotides. Additionally, we calculated the linear interaction energy between myosin and ADP.Mg^2+^ and observed no significant change in the interaction energy (WT: –347 ± 24; R403Q: –354 ± 24, *p =* 0.75). Our simulations suggest that modification of the myosin−thin filament interaction and/or detachment likely contributes more to the R403Q variant’s impaired relaxation than modification of myosin−ADP affinity.

## V. DISCUSSION

In this study, we employed a variety of biophysical, biochemical, and computational approaches to understand the detailed molecular mechanisms underlying altered contractile function in cardiac muscle containing the *MYH7* R403Q mutation, and to resolve apparently conflicting results in the literature (5–7). We used two model systems that naturally express predominantly *MYH7*: the Yucatan minipig model and a human induced pluripotent stem cell derived cardiomyocyte (hiPSC-CM) model, both of which contain the heterozygous R403Q mutation. Our results, consistent across both models, show that the mutation results in a hypercontractility at the tissue level under physiological Ca^2+^ levels typical of cardiac twitches, while slightly reducing the maximal force production capacity of the contractile organelles, i.e., myofibrils. The R403Q mutation also prolongs both the early, linear phase and major fast phase of relaxation. This appears to result at least in part from a reduced rate of cross-bridge detachment, as indicated by *k*_REL,slow_, which can prolong the initial phase of relaxation (t_REL,slow_). Computational simulations suggest this is not due to altered ADP release from the nucleotide-binding pocket, but rather by changes at the myosin-actin interface that retard cross-bridge dissociation. Finally, and importantly, we find that long-term cultured hiPSC-CMs can produce myofibrils with contractile properties similar to those of adult porcine myofibrils and replicate the effects of the R403Q mutation, similar to findings we recently reported for a cMyBP-C mutation that results in haploinsufficiency (47).

### The R403Q mutation increases the number and ATP turnover rate of myosins available for contraction in resting cardiac muscle

The greater submaximal force generated by R403Q myofibrils may result, at least in part, from a increased number of myosin heads that are initially available to bind to thin filaments once Ca^2+^ activates thin filaments. These additional myosin heads appear to come primarily from the thick filament D-zone, with some contribution from the P-zone. X-ray diffraction of relaxed minipig ventricular tissue suggests that the R403Q mutation disrupts the quasi-helical ordering of myosin heads on the thick filament backbone, similar to what occurs in contracting muscle, as indicated by reduced intensity of the first-order myosin-based layer line (I_MLL1_) and third-order myosin-based meridional reflection (I_M3_), and a significant increase in the sixth-order myosin-based meridional reflection (S_M6_). This greater disorder of myosin heads suggests a shift in the population of myosin heads from the OFF state toward a more ON state for R403Q cardiac muscle. Interestingly, we did not observe a change in the intensity ratio of the primary equatorial reflections (I_1,1_/I_1,0_), which is thought to be a measure of myosin head movement towards the thin filaments (17, 51). We have reported a similar pattern of change when dATP replaces ATP in relaxed cardiac muscle, but dATP also increases I_1,1_/I_1,0_ (18). One possible explanation for the R403Q results is that the stability of myosin on the thick filament backbone is disrupted by the mutation but the radial position of myosin relative to actin is not significantly affected, relative to WT tissue samples. Resting stiffness measurements of permeabilized cardiac strips support the notion that the R403Q mutation may not increase weak binding between actin and myosin (Supplemental Figure S3). In contrast, we have reported that dATP, a strong myosin activator, increases the positive charge on the actin binding surface (63) and increases electrostatic interactions between myosin and actin (52). This draws myosin heads toward thin filaments and increases weak binding. These dATP-induced electrostatic changes do not mimic the electrostatic landscape of myosin in the presence of the R403Q mutation. However, in relaxed R403Q muscle tissue where ATP was replaced with dATP, I_1,1_/I_1,0_ was significantly increased. The impact of dATP was less in R403Q porcine tissue compared to what we previously reported for WT porcine tissue (18), suggesting that the R403Q mutation has similar but incomplete effects on sarcomere structure compared to dATP. Like dATP, R403Q disrupts the IHM of myosin by the weakening of a salt-bridge interaction between blocked head R403 and free head Q454/Y455 (62, 64, 65). Our results from single-molecule tracking of ATP in isolated and relaxed myofibrils suggest that the R403Q mutation also increases the average ATPase activity of myosin heads, and overall shifts toward a smaller population of energy conserving, low ATPase heads. When subdivided into the P-, C-, and D-zones of the thick filament, this reduction occurs primarily in the P– and D-zones where cMyBP-C is not present. Although the distribution of R403Q myosin incorporation along the thick filament is unknown, cMyBP-C appears to be able to regulate the overall thick filament distribution of heads in R403Q myofibrils by moderately increasing the population of ON (higher ATPase) heads specifically in the C-zone, not due to changes in cMyBP-C phosphorylation (Supplemental Figure S5). Replacing the dark ATP nucleotide pool with dATP during single-molecule image acquisition demonstrated that dATP is a potent biochemical activator in WT myofibrils. The R403Q mutation had no further effect on the population of SRX heads. Although the validity of the mant-ATP chase experiment to quantify the SRX/DRX ratio remains under debate (66), our results from the single-molecule assay are consistent with reports using mant-ATP chase that suggest the R403Q mutation reduces the proportion of SRX heads in the R403Q minipig model (11), an R403Q rodent model (67), and for isolated 25-hep HMM constructs (64). Together, these structural and biochemical data suggest that the R403Q mutation results in both greater disorder of myosin heads at the thick filament backbone, as they shift toward more ON heads, but also elevated basal ATPase rates of these heads, priming myosins to be more available to bind to actin in the presence of Ca^2+^.

### Are the kinetics of altered myofibril contractile properties due to changes in actin-myosin interactions?

Reduced maximal force in R403Q porcine ventricular myofibrils is consistent with findings in myofibrils from human myectomy tissue (6, 7) and from the R403Q transgenic rabbit model (5). Optical trap experiments on isolated β-cardiac S1 have suggested reduced intrinsic and ensemble force (9). This suggests the R403Q mutation may reduce maximal myofibril force in part because increased myosin head recruitment does not fully compensate for diminished intrinsic force produced per myosin molecule.

While the R403Q mutation resulted in diminished maximal myofibril force consistently in porcine, human, and rabbit tissue, the slower contraction kinetics align with findings from the rabbit model but not with data from human myectomy tissue. Our CRISPR/Cas9-generated heterozygous R403Q hiPSC-CM model did not exhibit changes in activation kinetics or maximal force of myofibrils, but demonstrated impairments in relaxation consistent with the porcine R403Q myofibrils (Supplemental Figure S6). These differences in kinetics are not due to altered α/β-MHC ratios (Supplemental Figure S7), but may point to the necessity to control for confounding variables such as age, disease progression, and drug treatment history associated with HCM patients. Based on the slower rates of cross-bridge detachment during the isometric, slow phase of relaxation, this may result from impaired product release during the cross-bridge cycle. No changes in phosphate release were detected in the porcine R403Q tissue relative to WT (Supplemental Figure S8) and only modest changes in ADP release in the porcine tissue (but not in hiPSC-CMs) were identified (Supplemental Figure S9). Our MD simulations suggest a series of structural changes to myosin-ADP in its interaction with actin precede detachment in the force-bearing, post-powerstroke cross-bridge state. R403Q simulations showed increased contact frequency time between actin and myosin at multiple interfaces, while contact frequencies remained unimpaired within the nucleotide binding pocket between R403Q myosin and ADP.Mg^2+^. These structural changes at the actomyosin interface may be associated with changes in myosin detachment kinetics. The increased contact time at the actomyosin interface could also be associated with slower cross-bridge detachment (22).

Experiments performed in permeabilized tissue and myofibril preparations focused on the impact of the R403Q mutation on sarcomere function. To more closely mimic what occurs in the human heart, we compared WT and heterozygous R403Q hiPSC-CMs cast into 3D EHT constructs. Isometric twitch measurements at physiological, submaximal calcium (pCa 5.6) showed elevated peak tension in R403Q EHTs despite having smaller cross-sectional area. This is consistent with our findings of elevated Ca^2+^ sensitivity (pCa_50_) in the R403Q minipig permeabilized cardiac strips. Interestingly, this occurs despite reduced maximal force in isolated myofibrils.

In conclusion, with the two R403Q model systems presented, we have performed a multi-scale interrogation of the mechanisms of contractile dysfunction caused by the β-MHC R403Q mutation. With the aim of clarifying how the R403Q mutation affects contractile function of myofibrils, we show that the kinetics of contraction is slower than control myofibrils in young pigs and relaxation is slower than control myofibrils in young pigs and early stage HCM hiPSC-CMs. We suggest that instead of altered nucleotide handling, this mutation in the cardiomyopathy loop increases the binding interaction time between myosin and actin which may alter cross-bridge detachment speed despite there being enhanced structural and biochemical thick filament myosin recruitment in sub-sarcomere zones that do not contain cMyBP-C as determined by X-ray diffraction and single-molecule imaging.

## ACKNOWLEDGMENTS

The authors would like to thank Dr. Thomas Irving and Dr. Corrado Poggesi for their active discussions on this project. The authors acknowledge the Institute for Stem Cell and Regenerative Medicine’s Ellison Stem Cell Core at the University of Washington, as well as Dr. Nathan Sniadecki and Alex Goldstein (University of Washington) for their technical assistance with engineered heart tissues.

## AUTHOR CONTRIBUTIONS

MR, NK, FM-H, WM, and MG conceived and designed the research. SS, SM, MCC, TSM, AN, MD, KBK, CM, KT, JH, JM, and WM performed the experiments. SS, SM, MCC, TSM, MD, KT, JZ, JM, and WM analyzed the experimental data. SS, SM, MCC, TSM, AN, MP, MD, KBK, CM, JM, JMP, MAG, WM, FM-H, NK, and MR interpreted results of the experiments. SS, SM, MCC, TSM, and JM prepared the figures. SS and MR drafted the manuscript. All authors contributed to editing and revising the manuscript. All authors approved the final version for publication.

## SUPPLEMENTAL MATERIAL

**Supplemental Table S1.**
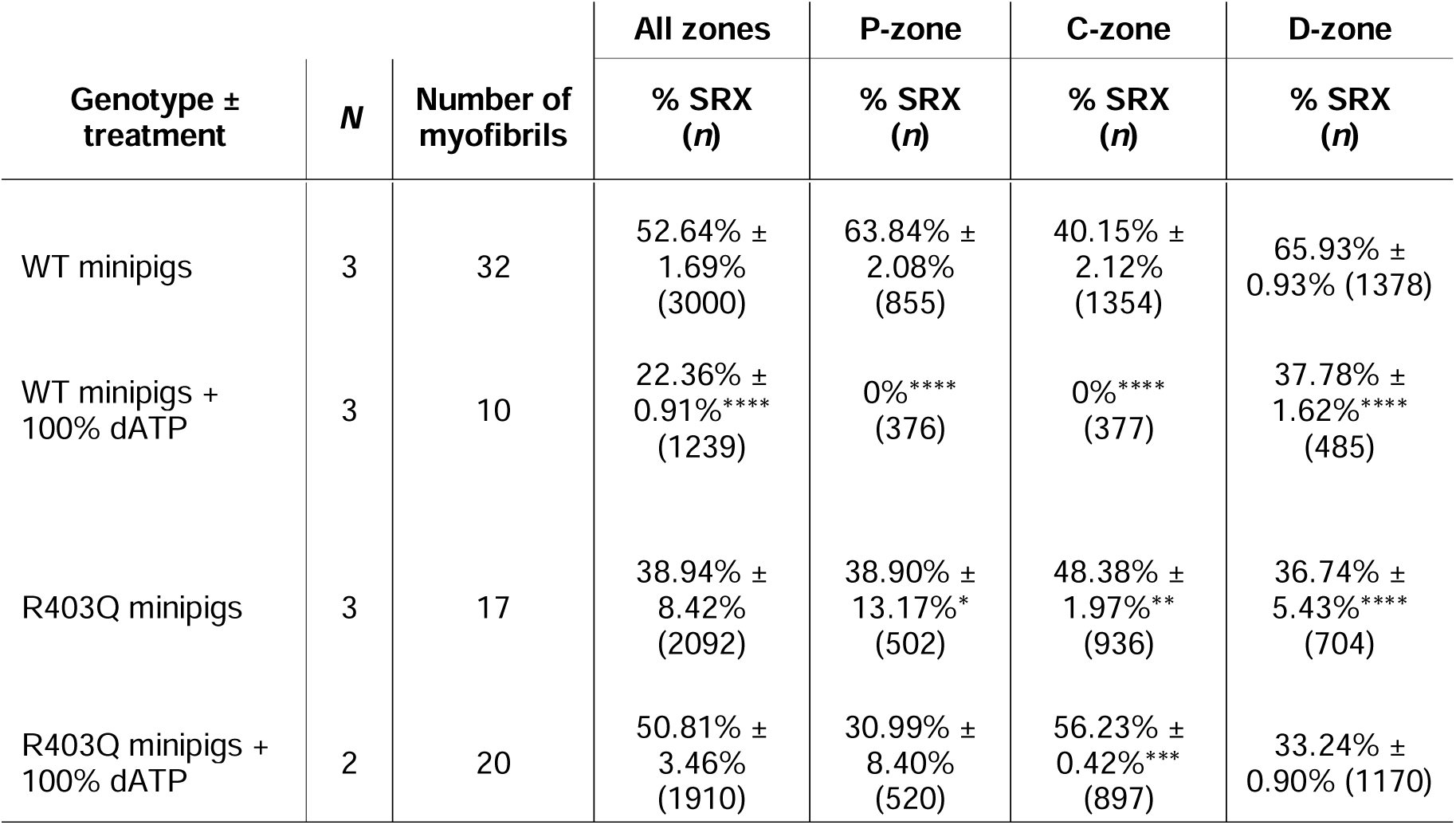
Quantification of the percentage of myosins in the SRX state in WT and R403Q porcine ventricular myofibrils. ATP binding lifetimes per ATP binding events were calculated and localized within the sub-sarcomere regions of the thick filament for WT and R403Q myofibrils. Single-molecule analysis was performed in the presence of saturating ATP or saturating dATP nucleotide (100%). Percentage super relaxed state (% SRX) myosin was calculated as an average across the entire sarcomere (‘all zones’) and sub-divided between the P-, C– and D-zones. *N* = number of minipig biological replicates. Number of events (*n =*) are provided for total tracked events and number of events attributed to each zone. Data are presented as mean ± SEM. Significance calculated with Student’s t-test, \**p <* 0.05, \*\**p <* 0.01, *** *p <* 0.001, **** *p <* 0.0001.

| Genotype ± treatment | N | Number of myofibrils | All zones | P-zone | C-zone | D-zone |
| --- | --- | --- | --- | --- | --- | --- |
|  |  |  | % SRX (n) | % SRX (n) | % SRX (n) | % SRX (n) |
| WT minipigs | 3 | 32 | 52.64% ±<br>1.69%<br>(3000) | 63.84% ±<br>2.08%<br>(855) | 40.15% ±<br>2.12%<br>(1354) | 65.93% ±<br>0.93% (1378) |
| WT minipigs +<br>100% dATP | 3 | 10 | 22.36% ±<br>0.91%****<br>(1239) | 0%****<br>(376) | 0%****<br>(377) | 37.78% ±<br>1.62%****<br>(485) |
| R403Q minipigs | 3 | 17 | 38.94% ±<br>8.42%<br>(2092) | 38.90% ±<br>13.17%*<br>(502) | 48.38% ±<br>1.97%**<br>(936) | 36.74% ±<br>5.43%****<br>(704) |
| R403Q minipigs +<br>100% dATP | 2 | 20 | 50.81% ±<br>3.46%<br>(1910) | 30.99% ±<br>8.40%<br>(520) | 56.23% ±<br>0.42%***<br>(897) | 33.24% ±<br>0.90% (1170) |

**Supplemental Table S2.** Summary of Yucatan minipig and hiPSC-CM myofibril force and kinetics parameters at maximal calcium concentration in the presence and absence of ADP. Myofibrils were isolated from both porcine ventricular tissue sections and hiPSC-CMs. Myofibril contractile experiments were performed at maximal calcium (pCa 4.5) with either 5 mM ATP or with 50:50 ATP:ADP (2.5 mM of each nucleotide). *k*_ACT_ = rate of activation; *k*_REL,slow_ = rate of slow phase of relaxation; t_REL,slow_ = duration of the slow phase of relaxation; *k*_REL,fast_ = rate of fast phase of relaxation. Data presented as mean ± SEM. *N* = number of minipig/hiPSC-CM biological replicates; *n* = number of myofibril technical replicates.

| [ATP],<br>mM | [ADP],<br>mM | Myofibril<br>group | $N, n$ | Specific<br>Force | $k_{\text{ACT}}$ | $k_{\text{REL,slow}}$ | $t_{\text{REL,slow}}$ | $k_{\text{REL,fast}}$ |
| --- | --- | --- | --- | --- | --- | --- | --- | --- |
|  |  |  |  | mN/mm<sup>2</sup> | s<sup>-1</sup> | s<sup>-1</sup> | ms | s<sup>-1</sup> |
| 5.0 | 0.0 | Porcine WT | 3, 19 | 83.93 ±<br>2.95 | 1.43 ±<br>0.08 | 0.58 ±<br>0.05 | 69.53 ±<br>3.38 | 10.53 ±<br>0.80 |
| | | Porcine R403Q | 3, 22 | 64.31 ±<br>3.02 | 1.07 ±<br>0.06 | 0.31 ±<br>0.02 | 121.27 ±<br>3.97 | 4.19 ±<br>0.30 |
| | | hiPSC-CM WT | 7, 18 - 19 | 50.54 ±<br>1.52 | 0.81 ±<br>0.07 | 0.27 ±<br>0.03 | 124.05 ±<br>5.30 | 4.47 ±<br>0.45 |
| | | hiPSC-CM R403Q | 6, 16 - 17 | 48.43 ±<br>2.97 | 0.72 ±<br>0.04 | 0.21 ±<br>0.02 | 168.71 ±<br>8.74 | 2.69 ±<br>0.23 |
| 2.5 | 2.5 | Porcine WT | 3, 19 | 78.21 ±<br>2.95 | 0.76 ±<br>0.03 | 0.21 ±<br>0.02 | 186.79 ±<br>7.57 | 3.41 ±<br>0.32 |
| | | Porcine R403Q | 3, 22 | 54.98 ±<br>2.64 | 0.60 ±<br>0.03 | 0.14 ±<br>0.01 | 274.73 ±<br>5.71 | 1.53 ±<br>0.09 |
| | | hiPSC-CM WT | 7, 18 - 19 | 45.53 ±<br>1.43 | 0.50 ±<br>0.03 | 0.14 ±<br>0.02 | 248.84 ±<br>12.18 | 2.21 ±<br>0.19 |
| | | hiPSC-CM R403Q | 6, 16 - 17 | 44.63 ±<br>3.66 | 0.48 ±<br>0.02 | 0.12 ±<br>0.02 | 303.59 ±<br>14.10 | 1.43 ±<br>0.11 |

**Supplemental Figure S1.**
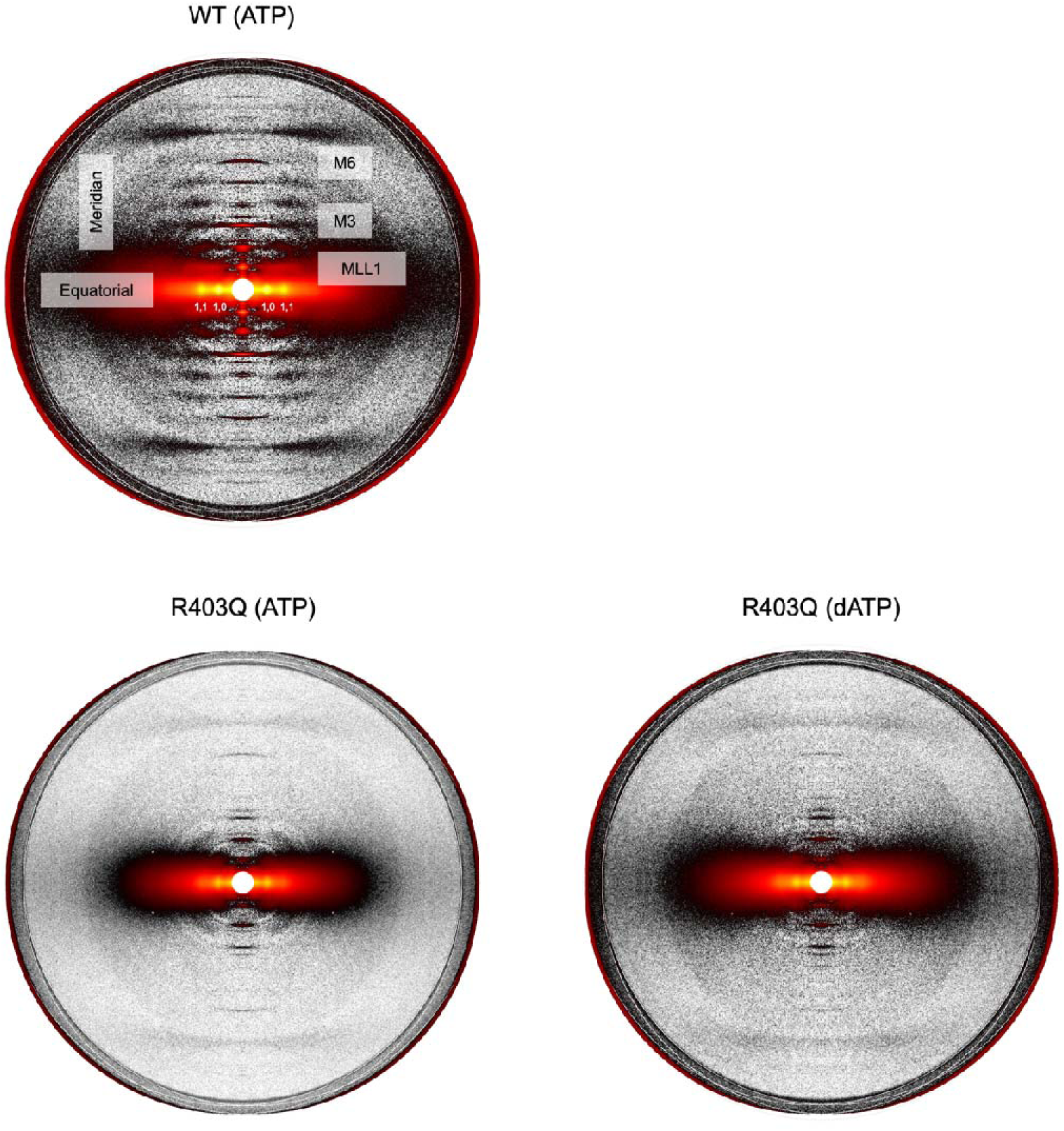
Original (un-cropped and un-edited), sample X-ray diffraction patterns used for quantification. Sample X-ray diffraction patterns of (A) WT and (B) R403Q porcine ventricular myocardium under resting conditions. (C) Sample X-ray diffraction pattern of R403Q myocardium incubated with 100% dATP at the time of data collection.

**Supplemental Figure S2.**
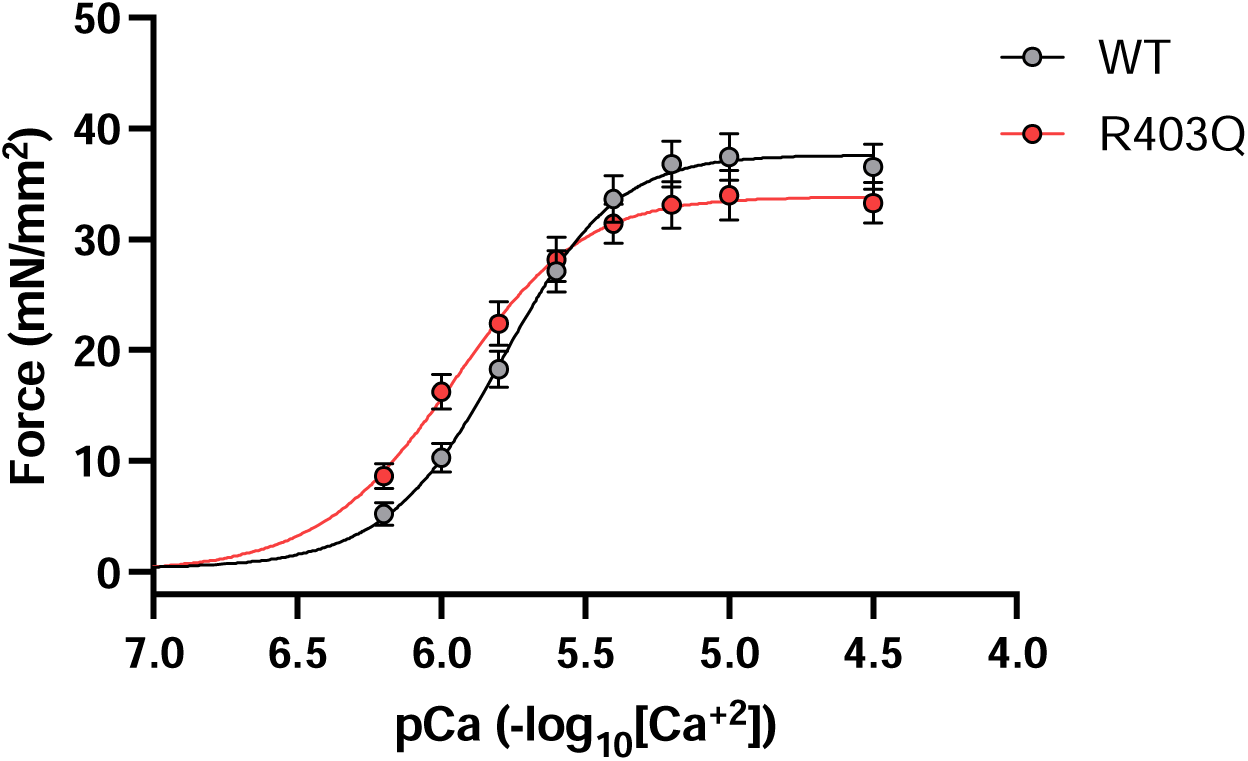
Force-pCa curves from isolated uniaxially aligned porcine ventricle tissue strips from WT (grey line) and R403Q (red line) littermate minipigs. WT: *N =* 3, *n =* 14; R403Q: *N =* 3, *n =* 14. Data are presented as mean ± SEM. *N =* number of biological replicate minipigs; *n =* number of technical replicates.

**Supplemental Figure S3.**
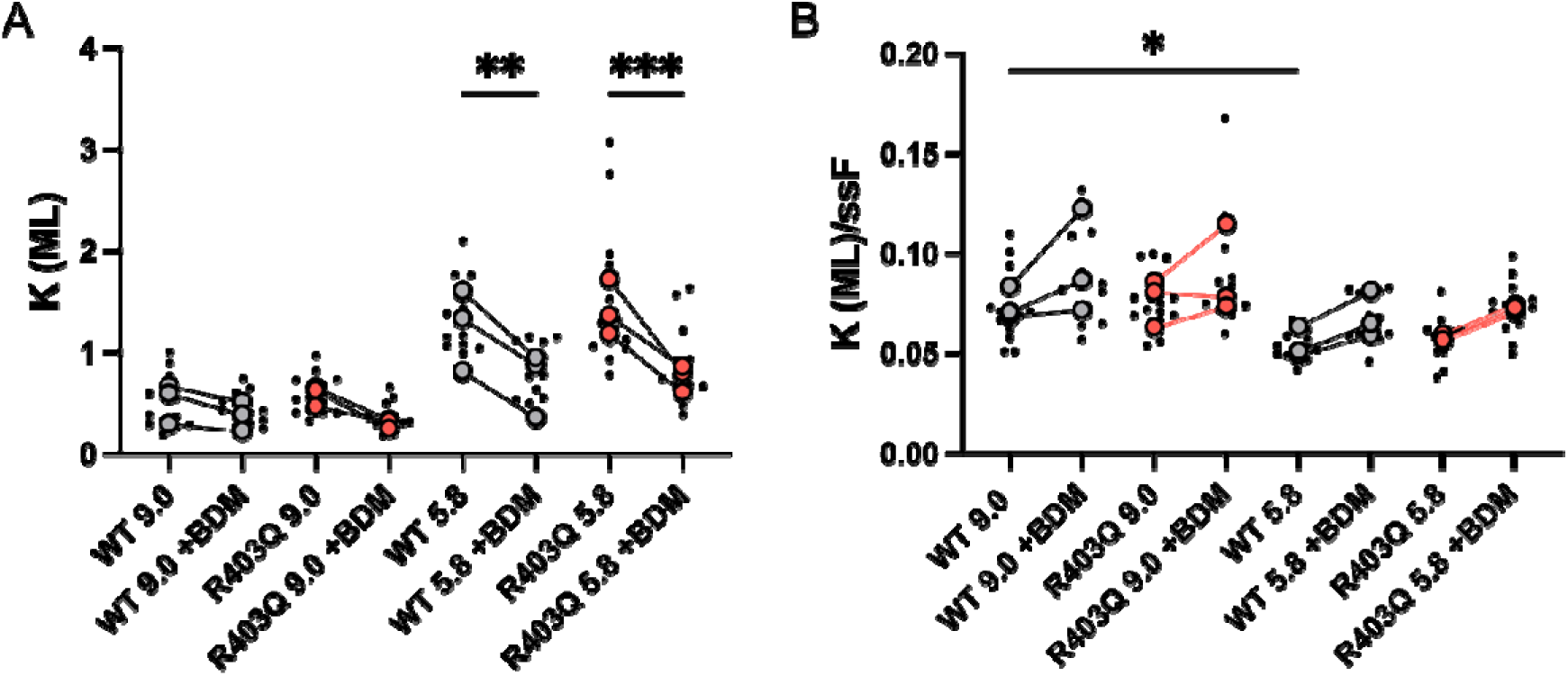
No differences in tissue stiffness were identified between WT (grey) and R403Q (red) myocardium. High-frequency stiffness (HFS) was measured by applying a 1000 Hz sinusoidal length change (±0.5% muscle length (ML)) under relaxed conditions and submaximal calcium activation (pCa 5.8). To exclude impact of bound cross-bridges, stiffness was also measured in the presence of BDM. Two-way ANOVA with Tukey post hoc test, \**p <* 0.05, \*\**p <* 0.01, \*\*\**p <* 0.001.

**Supplemental Figure S4.**
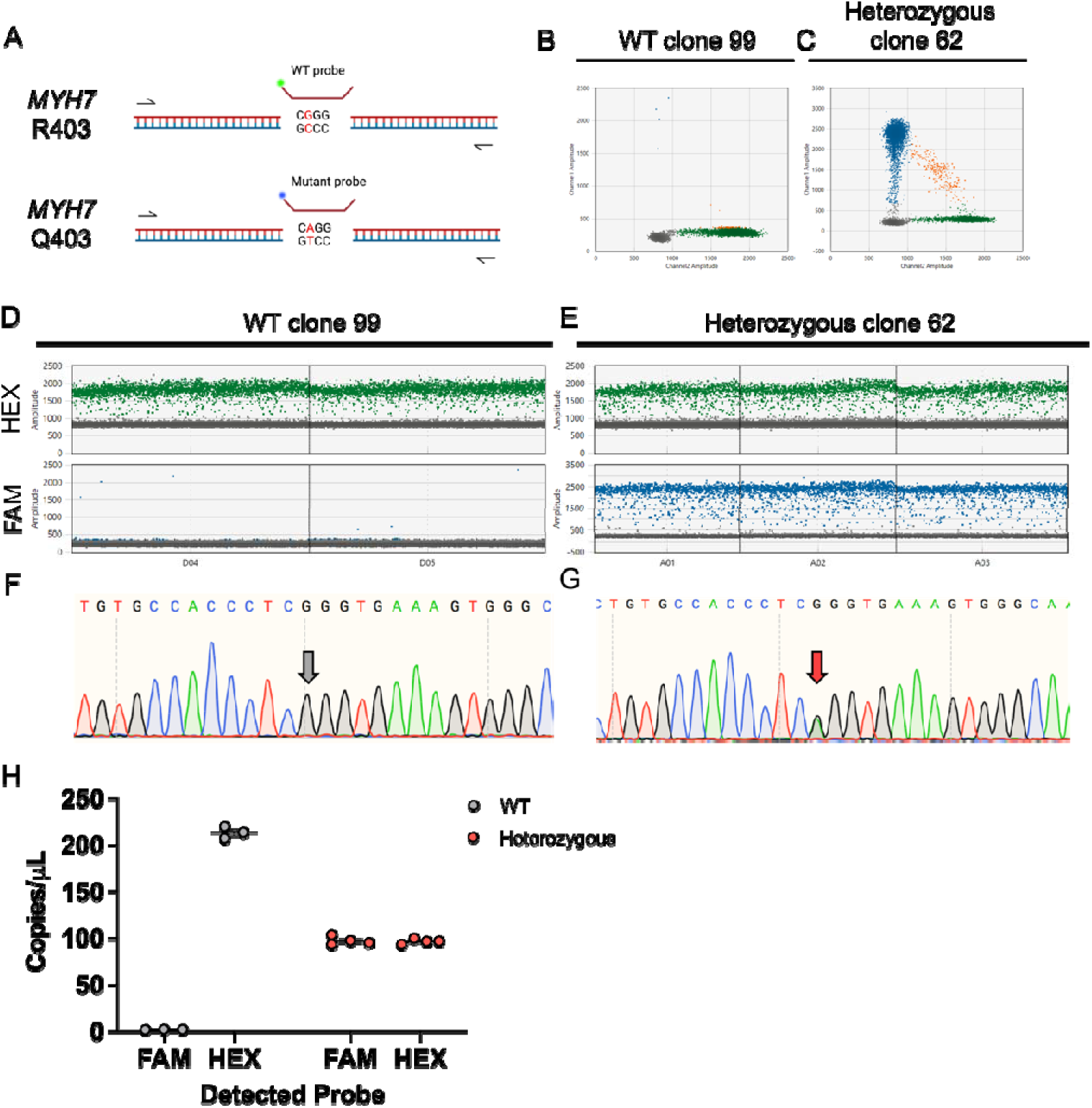
Digital droplet PCR (ddPCR) was used to confirm correct CRISPR/Cas9 genome editing to insert the expected *MYH7* c.1208G>A (R403Q) mutation into human, wildtype WTC-11 hiPSCs. (A) Schematic of ddPCR assay set up. A primer pair and two probes (a HEX-wildtype (WT) probe and a FAM-mutant probe) were used to detect both mutant and WT alleles in the same reaction. 2D amplitude view showing (B) WT clone 99 and (C) heterozygous clone 62. 1D amplitude view of HEX and FAM probes for (D) WT and (E) heterozygous. Accompanying Sanger Sequencing for (F) WT and (G) heterozygous clones. (H) ddPCR concentration graph showing the number of DNA copies per microliter detected with each probe (WT: *N =* 1, *n =* 3, heterozygous: *N =* 1, *n =* 4). *N =* number of biological replicates; *n =* number of technical replicates.

**Supplemental Figure S5.**
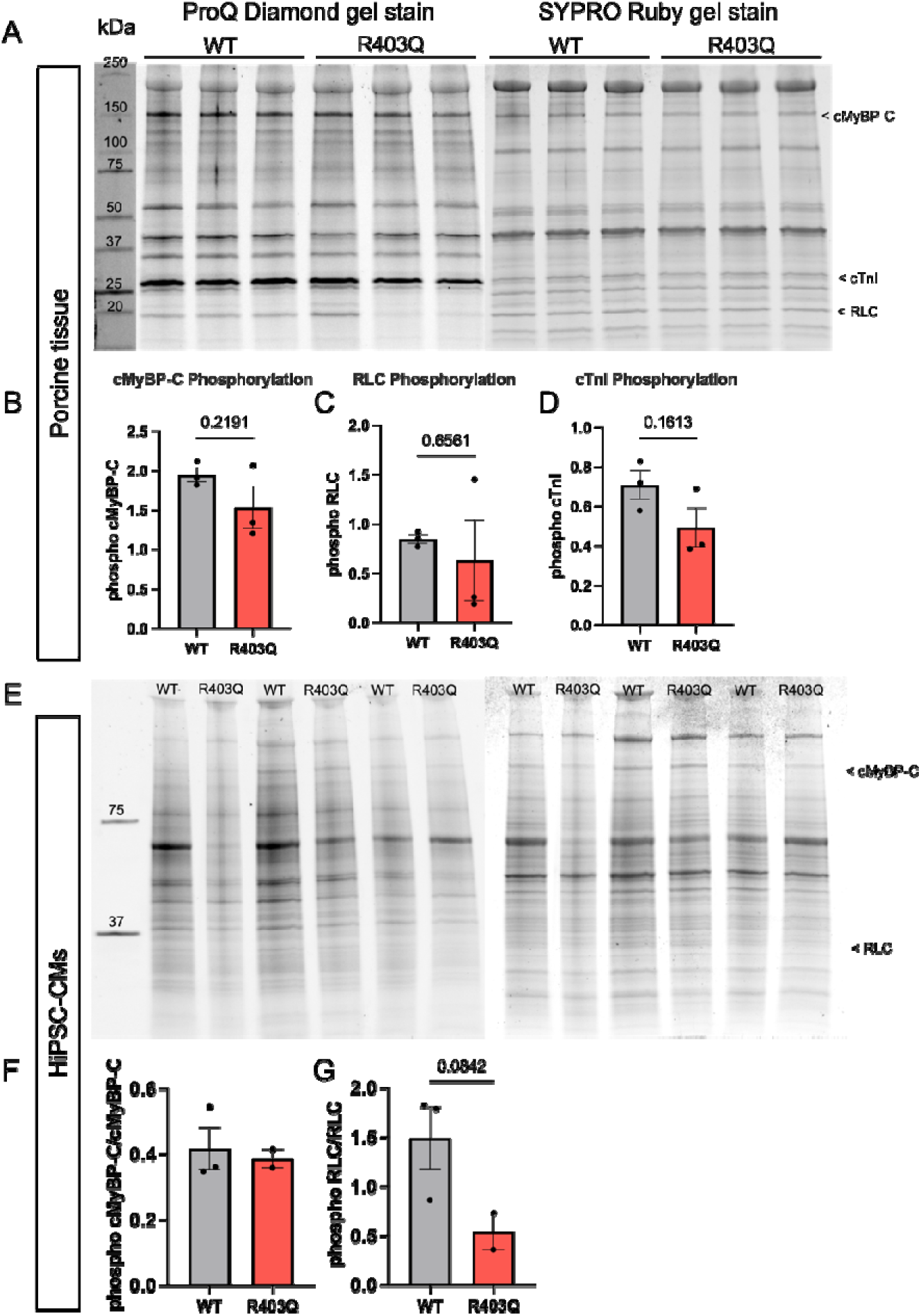
Phosphorylation profile of key sarcomeric proteins, cMyBP-C, RLC, and cTnI in both porcine myofibrils and human iPSC-CM cell lysates. (A) Uncropped and unedited Pro-Q Diamond and Coomassie total protein gel stains on lysates from WT and R403Q isolated porcine ventricular myofibrils. Densitometry analysis was performed on cMyBP-C, RLC, and cTnI bands from both gels, with Pro-Q phosphorylation bands normalized to corresponding total protein band. Quantification of phosphorylated (B) cMyBP-C, (C) RLC, and (D) cTnI (WT: *N =* 3, *n =* 1; R403Q: *N =* 3, *n =* 1). (E) Uncropped and unedited Pro-Q Diamond and Coomassie total protein gel stains on lysates from day 45 – 46 WT and heterozygous R403Q hiPSC-CMs. Quantification of phosphorylated (F) cMyBP-C and (G) RLC (WT: *N =* 3, *n =* 1; R403Q: *N =* 2, *n =* 1). No quantification of cTnI band was possible in hiPSC-CM samples. Ladder used was a Precision Plus Protein^™^ Dual Color Standards ladder. Data are presented as mean ± SEM. *N =* number of biological replicates; *n =* number of technical replicates run per sample. *P* value was determined by unpaired, two-tailed Student’s t-test with Welch’s correction.

**Supplemental Figure S6.**
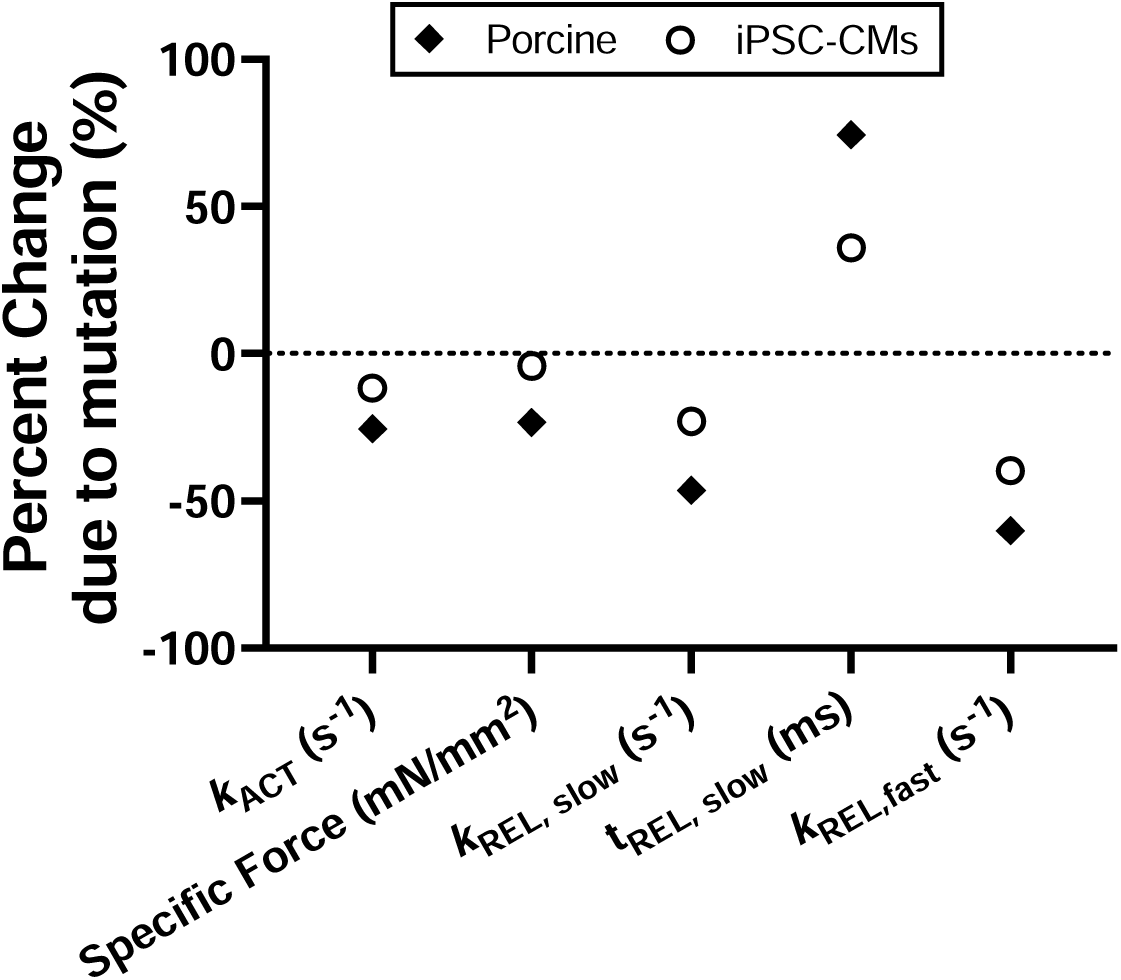
R403Q mutation induced changes in myofibril kinetics parameters exhibit similarities between day 45 hiPSC-CM derived myofibrils and porcine ventricular myofibrils. For each myofibril kinetics parameter calculated from the isolated myofibril experiment at maximal calcium activation (pCa 4.5), percent change was calculated between the mutant and its corresponding control. The dashed black line corresponds to zero, indicating no change between mutant and control.

**Supplemental Figure S7.**
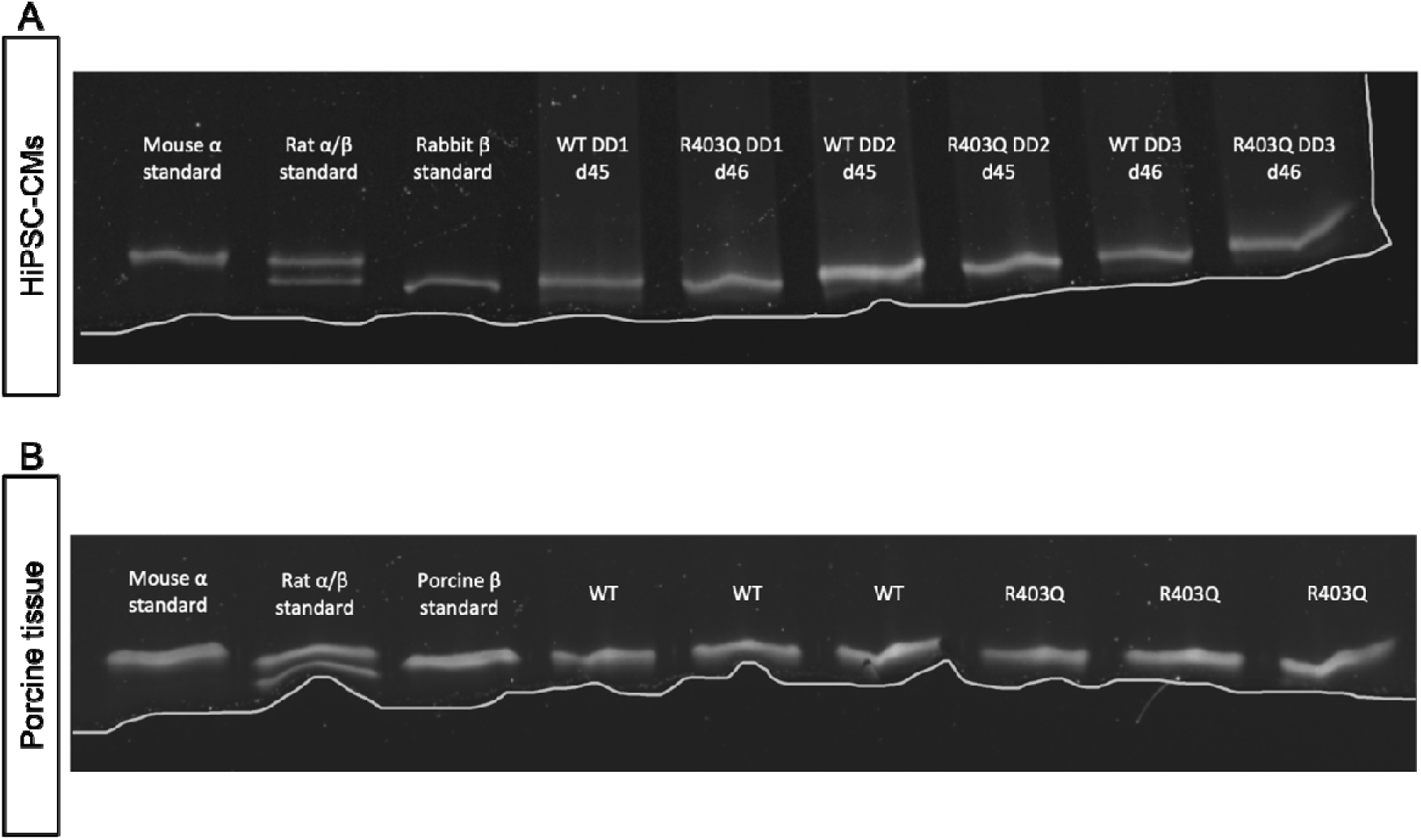
Gel separation between (A) human hiPSC-CM and (B) porcine α– and β-MHC isoforms for WT and heterozygous R403Q samples. For human MHC, α-MHC isoform is 224 kDa, while the β-MHC isoform is 223 kDa. (A) For hiPSC-CM samples, day 45 – 46 whole cell protein lysates were run on a myosin separation gel for WT (*N =* 3, *n =* 1) and R403Q (*N =* 3, *n =* 1) samples. Controls were protein lysates from isolated mouse, rat, and rabbit cardiac tissue myofibrils (top gel). (B) For porcine samples, proteins were lysed from isolated myofibrils and run on a separation gel for WT (*N =* 3, *n =* 1) and R403Q (*N =* 3, *n =* 1) samples. Controls were protein lysates from isolated mouse, rat and non-transgenic porcine cardiac tissue myofibrils (bottom gel). *N =* number of biological replicate minipigs; *n =* number of technical replicates. A white line was manually drawn to designate the border of the gel.

**Supplemental Figure S8.**
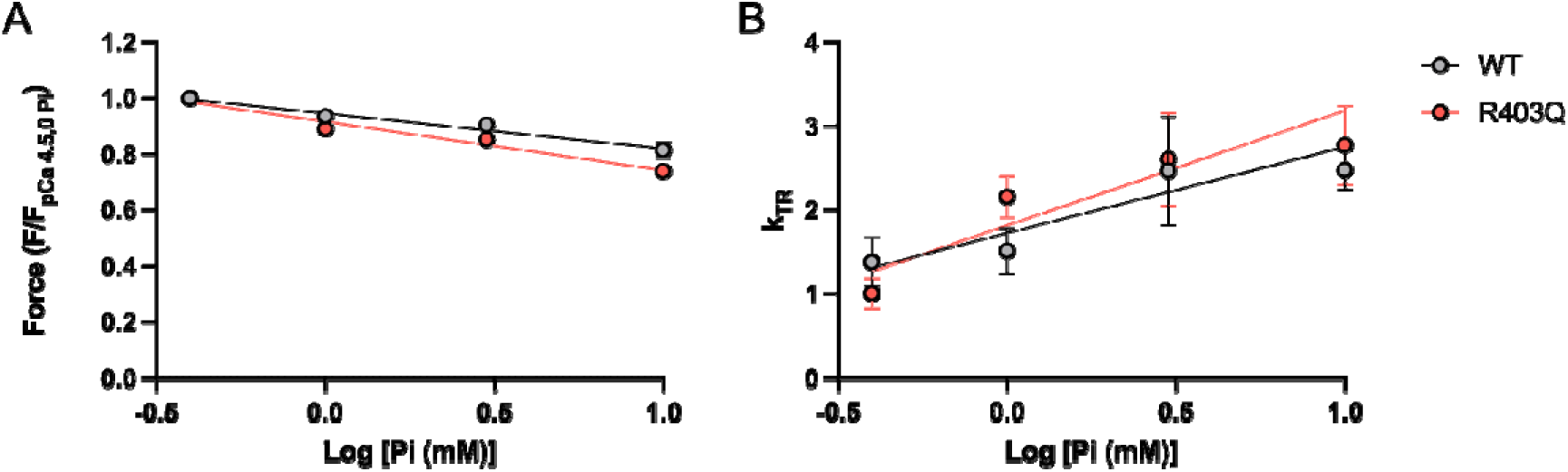
Permeabilized WT and R403Q ventricular strips did not exhibit impairments in phosphate release. (A) Force and (B) *k*_TR_ measurements for WT and R403Q porcine ventricular tissue were taken in the presence of increasing concentrations of inorganic phosphate (Pi). The linear relationship between relative force and log[Pi (mM)] in permeabilized mechanics was plotted for both parameters. Data are presented as mean ± SEM.

**Supplemental Figure S9.**
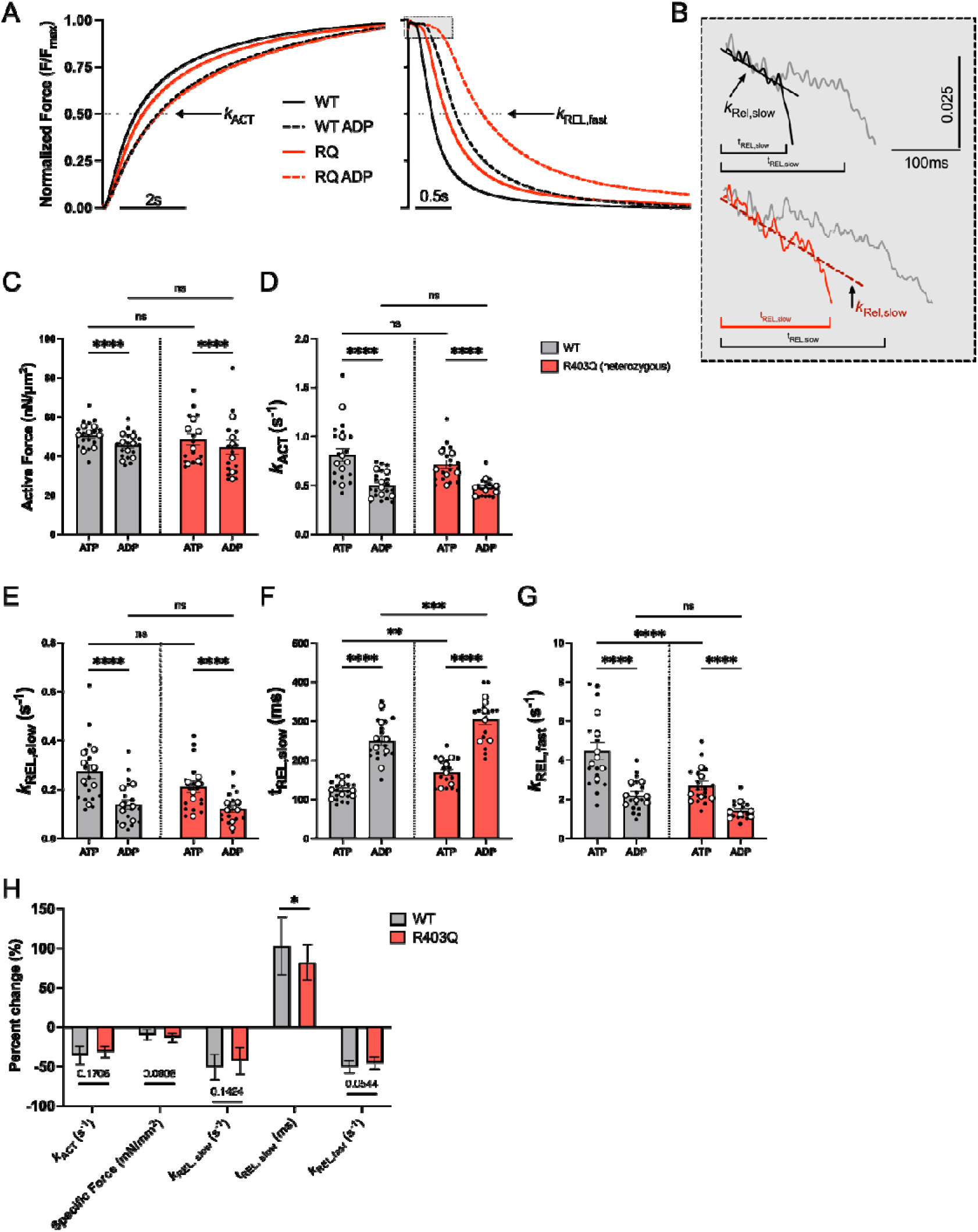
HiPSC-CM myofibrils with the R403Q mutation exhibit less alteration of ADP release than adult myofibrils. (A) Average normalized myofibril kinetics traces for WT and R403Q hiPSC-CM myofibrils with and without the presence of elevated ADP in the maximal calcium activation solution (pCa 4.5). (B) Traces highlighting the slow phase of relaxation (with a manual offset for visualization) before and after ADP inhibition experiment for WT (top black: 100% ATP, top grey: 50% ATP:50% ADP) and R403Q (bottom red: 100% ATP, bottom grey: 50% ATP:50% ADP) hiPSC-CM lines. Quantification of (C) maximal force, (D) rate of activation (E) rate of the slow phase of relaxation, (F) duration of the slow phase of relaxation, and (G) rate of the fast phase of relaxation before and after paired ADP inhibition experiment (WT | ATP: *N =* 7, *n =* 19; WT | ADP: *N =* 7, *n =* 18 – 19; R403Q | ATP: *N =* 6, *n =* 17, R403Q | ADP: *N =* 6, *n =* 16 – 17). (H) Percent change in each myofibril parameter after ADP inhibition experiment for both WT and R403Q hiPSC-CMs. Data presented as mean ± SEM. *N =* number of biological replicate differentiation batches; *n =* number of myofibril technical replicates. Two-way ANOVA with Tukey post-hoc test, \**p <* 0.05, \*\**p <* 0.01, \*\*\**p <* 0.001, \*\*\*\**p <* 0.0001.

